# Assessment of human diploid genome assembly with 10x Linked-Reads data

**DOI:** 10.1101/729608

**Authors:** Lu Zhang, Xin Zhou, Ziming Weng, Arend Sidow

## Abstract

**Background:** Producing cost-effective haplotype-resolved personal genomes remains challenging. 10x Linked-Read sequencing, with its high base quality and long-range information, has been demonstrated to facilitate *de novo* assembly of human genomes and variant detection. In this study, we investigate in depth how the parameter space of 10x library preparation and sequencing affects assembly quality, on the basis of both simulated and real libraries.

**Findings:** We prepared and sequenced eight 10x libraries with a diverse set of parameters from standard cell lines NA12878 and NA24385 and performed whole genome assembly on the data. We also developed the simulator LRTK-SIM to follow the workflow of 10x data generation and produce realistic simulated Linked-Read data sets. We found that assembly quality could be improved by increasing the total sequencing coverage (*C*) and keeping physical coverage of DNA fragments (*C_F_*) or read coverage per fragment (*C_R_*) within broad ranges. The optimal physical coverage was between 332X and 823X and assembly quality worsened if it increased to greater than 1,000X for a given *C*. Long DNA fragments could significantly extend phase blocks, but decreased contig contiguity. The optimal length-weighted fragment length (*Wμ_FL_*) was around 50 – 150kb. When broadly optimal parameters were used for library preparation and sequencing, ca. 80% of the genome was assembled in a diploid state.

**Conclusion:** The Linked-Read libraries we generated and the parameter space we identified provide theoretical considerations and practical guidelines for personal genome assemblies based on 10x Linked-Read sequencing.

## Data description

### Introduction

The human genome holds the key for understanding the genetic basis of human evolution, hereditary illnesses and many phenotypes. Whole-genome reconstruction and variant discovery, accomplished by analysis of data from whole-genome sequencing experiments, are foundational for the study of human genomic variation and analysis of genotype-phenotype relationships. Over the past decades, cost-effective whole-genome sequencing has been revolutionized by short-fragment approaches, the most widespread of which have been the consistently improving generations of the original Solexa technology [1, 2], now referred to as Illumina sequencing. Illumina’s strengths and weaknesses are inherent in the sample preparation and sequencing chemistry. Illumina generates short paired reads (2×150 base pairs for the highest-throughput platforms) from short fragments (usually 400-500 base pairs) [3]. Because many clonally amplified molecules generate a robust signal during the sequencing reaction, Illumina’s average per-base error rates are very low.

The lack of long-range contiguity between end-sequenced short fragments limits their application for reconstructing personal genomes. Long-range contiguity is important for phasing variants and dealing with genomic complex regions. For haplotyping, variants can be phased by population-based methods [4, 5] or family-based recombination inference [6, 7]. However, such approaches are only feasible for common variants in single individuals or when a trio or larger pedigree is sequenced. Furthermore, highly polymorphic regions such as the HLA in which the reference sequence does not adequately capture the diversity segregating in the population are refractory to mapping-based approaches and require *de novo* assembly to reconstruct [8]. Short-read/short-fragment data are challenged by interspersed repetitive sequences from mobile elements and by segmental duplications, and only support highly fragmented genome reconstruction [9, 10].

In principle, many of these challenges can be overcome by long-read/long-fragment sequencing [11, 12]. Assembly of Pacific Biosciences (PacBio) or Oxford Nanopore (ONT) data can yield impressive contiguity of contigs and scaffolds. In one study [13], scaffold N50 reached 31.1Mb by hierarchically integrating PacBio long reads and BioNano for a hybrid assembly, which also uncovered novel tandem repeats and replicated the structural variants that were newly included in the updated hg38 human reference sequence. Another study [14] produced human genome assemblies with ONT data, in which a contig N50 ~3Mb was achieved, and long contigs covered all class I HLA regions. A recent whole genome assembly of NA24385 [15] with high quality PacBio CCS reads generated contigs with an N50 of 15Mb. However, long-fragment sequencing suffers from extremely high cost (in the case of PacBio CCS), or low base quality (in the case of single-pass reads of either technology), hampering its usefulness for personal genome assembly.

Hierarchical assembly pipelines in which multiple data types are used as another approach for genome assembly [16]. For example, in the reconstruction of an Asian personal genome, fosmid clone pools and Illumina data were merged, but because fosmid libraries are highly labor intensive to generate and sequence, this approach is not generalizable to personal genomes. The “Long Fragment Read” (LFR) approach [17], where a long fragment is sequenced at high depth via single-molecule fragmented amplification, reported promising personal genome assembly and variant phasing by attaching a barcode to the short reads derived from the same long fragment. However, because LFR is implemented in a 384 well plate, many long fragments would be labelled by the same barcodes, making it difficult for binning short-reads, and the great sequencing depth required rendered LFR not cost-effective.

An alternative approach is offered by the 10x Genomics Chromium system, which distributes the DNA preparation into millions of partitions where partition-specific barcode sequences are attached to short amplification products that are templated off the input fragments. Because of the limited reaction efficiency in each partition, the sequencing depth for each fragment is too shallow to reconstruct the original long-fragment, distinguishing this approach from LFR [18]. However, to compensate for the low read coverage of each fragment, each genomic region is covered by hundreds of DNA fragments, giving overall sequence coverage that is in a range comparable to standard Illumina short-fragment sequencing while providing very high physical coverage. Novel computational approaches leveraging the special characteristics of 10x Genomics data have already generated significant advances in power and accuracy of haplotyping [19, 20], cancer genome reconstruction [21, 22], metagenomic assemblies [23], and *de novo* assembly of human and other genomes [24–26], compared to standard Illumina short-fragment sequencing. While the uniformity of sequence coverage is not as good as with PCR-free Illumina libraries, 10x Linked-Read sequencing is a promising technology that combines low per-base error and good small-variant discovery with long-range information for much improved SV detection in mapping-based approaches [22, 27], and the possibility of long-range contiguity in *de novo* assembly [24, 26, 28].

Practical advantages of the technology include the low DNA input mass requirement (1ng per library, or approximately 300 haploid human genome equivalents). Real input quantities can vary, along with other factors, to influence an interconnected array of parameters that are relevant to genome assembly and reconstruction. The parameters over which the experimenter has influence are (**Figure 1**): i). *C_R_*: average **<u>C</u>**overage of short **<u>R</u>**eads per fragment; ii). *C_F_*: average physical **<u>C</u>**overage of the genome by long DNA **<u>F</u>**ragments; iii). *N_F/P_*: **<u>N</u>**umber of **<u>F</u>**ragments per **<u>P</u>**artition; iv). Fragment length distribution, several parameters of which are used, specifically *μ_FL_*: Average Unweighted DNA **<u>F</u>**ragment **<u>L</u>**ength and W*μ_FL_*: Length-**<u>W</u>**eighted average of DNA **<u>F</u>**ragment **<u>L</u>**ength. Note that several parameters depend on each other. For example, a greater amount of input DNA will increase *N_F/P_*; shorter fragments increase *N_F/P_* at the same DNA input amount compared to longer fragments; less input DNA will (within practical constraints) increase *C_R_* and decrease *C_F_*; and their absolute values are set by how much total sequence coverage is generated because *C_R_* x *C_F_* = *C*.

**Figure 1.**
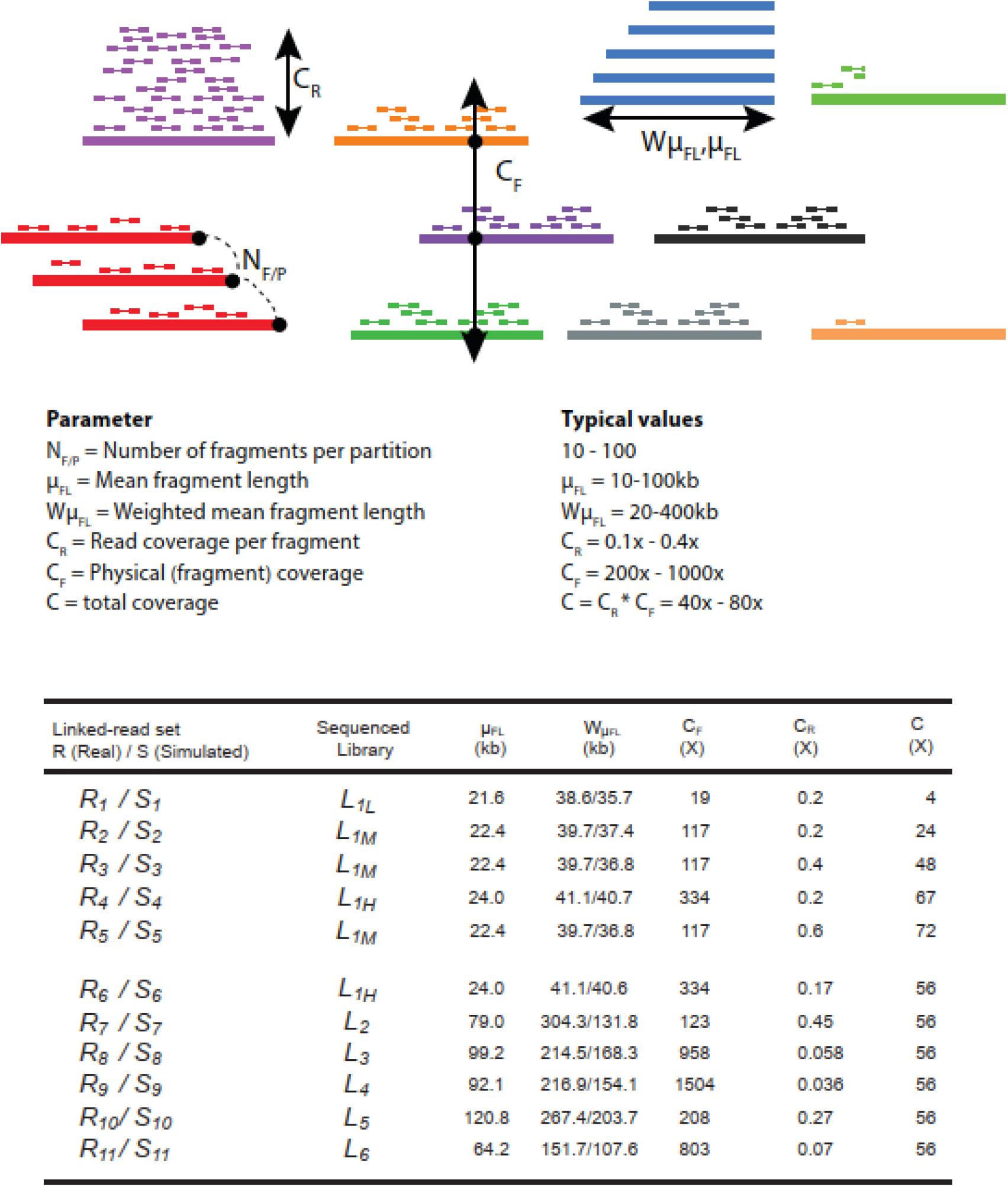
The linked-read sets prepared to evaluate the impact of *C_F_, C_R_, μ_FL_* and *Wμ_FL_* on human diploid assembly.

Our goal in this study was to experimentally explore the 10x parameter space and evaluate the quality of *de novo* diploid assembly as a function of the parameter values. For example, we set out to ask whether longer input fragments produce better assemblies, or what the effect of sequencing vs. physical coverage is on contiguity of assembly. In order to constrain the parameter space, we first performed computer simulations with reasonably realistic synthetic data. The simulation results suggested certain parameter combinations that we then approximated in the generation of real, high-depth, sequence data on two human reference genome cell lines, NA12878 and NA24385. These simulated and real data sets were then used to produce *de novo* assemblies, with an emphasis on the performance of 10x’s Supernova2 [24]. We finally assessed the quality of the assemblies using standard metrics of contiguity and accuracy, facilitated by the existence of a gold standard (in the case of simulations) and comparisons to the reference genome (in the case of real data).

### Library preparation, physical parameters and sequencing coverage

We made six DNA preparations that varied in fragment size distribution and amount of input DNA, three each from NA12878 and NA24385. From these, we prepared eight libraries, five from NA12878 and three from NA24385 (**Table S1**). To generate libraries *L*_1*L*_, *L*_1*M*_ and *L*_1*H*_ (the subscripts *L, M* and *H* represent low, medium and high C_F_, respectively), genomic DNA was extracted from ca. 1 million cultured NA12878 cells using the Gentra Puregene Blood Kit following manufacturer’s instructions (Qiagen, Cat. No 158467). The GEMs were divided into 3 tubes with 5%, 20%, and 75% to generate libraries *L*_1*L*_, *L*_1*M*_ and *L*_1*H*_, respectively (**Figure S1-S3**). For the other libraries, to generate longer DNA fragments (W*μ_FL_*=150kb and longer, **Figure S4-S8**), a modified protocol was applied. Two-hundred thousand NA12878 or NA24385 cells of fresh culture were added to 1mL cold 1x PBS in a 1.5 ml tube and pelleted for 5 minutes at 300g. The cell pellets were completely resuspended in the residual supernatant by vortexing and then lysed by adding 200ul Cell Lysis Solution and 1ul of RNaseA Solution (Qiagen, Cat. No 158467), mixing by gentle inversion, and incubating at 37°C for 15-30 minutes. This cell lysis solution is used immediately as input for the 10x Chromium preparation (ChromiumTM Genome Library & Gel Bead Kit v2, PN-120258; ChromiumTM i7 Multiplex Kit, PN-120262). Fragment size of the input DNA can be controlled by gentle handling during lysis and DNA preparation for Chromium. The amount of input DNA (between 1.25 and 4 ng) was varied to achieve a wide range of physical coverage (*C_F_*).The Chromium Controller was operated and the GEM preparation was performed as instructed by the manufacturer. Individual libraries were then constructed by end repairing, A-tailing, adapter ligation and PCR amplification. All libraries were sequenced with three lanes of paired-end 150bp runs on the Illumina HiSeqX to obtain very high coverage (*C*=94x-192x), though the two with the fewest number of gel beads (*L*_1*L*_ and *L*_1*M*_) exhibited high PCR duplication rates because of the reduced complexity of the libraries (**Table S1**).

### Linked-Reads subsampling

The high sequencing coverage in the libraries allowed subsampling to facilitate the matching of parameters among the different libraries, for purposes of comparability; these subsampled Linked-Read sets are denoted *R_id_* (**Figure 1**). We aligned the 10x Linked-Reads to human reference genome (hg38, GRCh38 Reference 2.1.0 from 10x website) followed by removing PCR duplication by barcode-aware analysis in Long Ranger[21]. Original input DNA fragments were inferred by collecting the read-pairs with the same barcode that were aligned in proximity to each other. A fragment was terminated if the distance between two consecutive reads with the identical barcode larger than 50kb. Fragments were required to have at least two read pairs with the same barcode and a length of at least 2 kb. Partitions with fewer than three fragments were removed. We subsampled short-reads for each fragment to satisfy the expected *C_R_*.

### Generating 10x simulated libraries by LRTK-SIM

To compare the observations from real data with a known truth set, we developed LRTK-SIM, a simulator that follows the workflow of the 10x Chromium system and generates synthetic Linked-Reads like those produced by an Illumina HiSeqX machine (**Supplementary Information** and **Figure S9**). Based on the parameters commonly employed by 10x Genomics Linked-Read sequencing and the characteristics of our libraries, LRTK-SIM generated simulated datasets from the human reference (hg38), explicitly modeling the five key steps in real data generation. Parameters in parentheses are from the standard 10x Genomics protocol: 1. Shearing genomic DNA into long fragments (W*μ_FL_* from 50kb to 100kb); 2. Loading DNA to the 10x Chromium instrument (~1.25ng DNA); 3. Allocating DNA fragments into partitions which are attached the unique barcodes (~10 fragments per partition); 4. Generating short fragments; 5. Generating Illumina paired-end short reads (800M~1200M reads). LRTK-SIM first generated a diploid reference genome as a template by duplicating the human reference genome (hg38) into two haplotypes and inserting SNVs from high-confidence regions in GIAB of NA12878 (ftp://ftp-trace.ncbi.nlm.nih.gov/giab/ftp/release/NA12878_HG001/latest/GRCh38/HG001_GRCh38_GIAB_highconf_CG-IllFB-IllGATKHC-Ion-10X-SOLID_CHROM1-X_v.3.3.2_highconf_nosomaticdel_noCENorHET7.bed); For low-confidence regions we randomly simulated 1 SNV per 1 kb. The ratio was 2:1 for heterozygous and homozygous SNVs. From this diploid reference genome, LRTK-SIM generated long DNA fragments by randomly shearing each haplotype with multiple copies into pieces whose lengths were sampled from an exponential distribution with mean of *μ_FL_*. These fragments were then allocated to pseudo-partitions, and all the fragments within each partition were assigned the same barcode. The number of fragments for each partition was randomly picked from a Poisson distribution with mean of *N_F/P_*. Finally, paired-end short reads were generated according to *C_R_* and replaced the first 16bp of the reads from forward strand to the assigned barcodes followed by 7 Ns. More information about implementation can be found in Supplementary Information. From that diploid genome, Linked-Read datasets were generated that varied in *C_R_*, *C_F_* and (W*μ_FL_*) (**Table S2-S3**). Varying *N_F/P_* was only done for chromosome 19 because of the infeasibility of running Supernova2 on whole genome assemblies with large *N_F/P_*; within practically reasonable values, *N_F/P_* does not appear to influence assembly quality (**Figure S10**). In total, we generated 17 simulated Linked-Read datasets to explore the overall parameter space (**Table S2-S3**) and 11 to match the parameters of the abovementioned real libraries (**Figure 1**).

### Human genome diploid assembly and evaluation

The scaffolds were generated by the “pseudohap2” output of Supernova2, which explicitly generated two haploid scaffolds, simultaneously. Contigs were generated by breaking the scaffolds if at least 10 consecutive ‘N’s appeared, per definition by Supernova2. For the simulations of human chromosome 19, we used the scaffolds from the “megabubbles” output. Contig and scaffold N50 and NA50 were used to evaluate assembly quality. Contigs longer than 500bp were aligned to hg38 by Minimap2[29]. We calculated contig NA50 on the basis of contig misassemblies reported by QUAST-LG [30]. For scaffolds (longer than 1kb), we calculated the NA50 following Assemblathon 1’s procedure [31] (**Supplementary Information**).

### Genomic variant calls from diploid assembly

We compared single nucleotide variants (SNVs) and structural variants (SVs) from the diploid regions of our assemblies with the ones from standard Illumina data and reference-based processing of our 10x data. The standard Illumina data were downloaded from Genome in a Bottle [32] and analyzed with SVABA [33] to generate SV calls, and with BWA [34] and FreeBayes [35] to generate SNV calls. Long ranger (https://support.10xgenomics.com/genome-exome/software/pipelines/latest/what-is-long-ranger) was used to generate SNV and SV (only deletions) calls for 10x reference-based analysis. We noted that R_9_ failed to be analyzed by Long Ranger due to its extremely large C_F_. For SNVs, we benchmarked the calls from three strategies using the gold standard of NA12878 (ftp://ftp-trace.ncbi.nlm.nih.gov/giab/ftp/release/NA12878_HG001/latest/GRCh38/) and NA24385 (ftp://ftp-trace.ncbi.nlm.nih.gov/giab/ftp/release/AshkenazimTrio/HG002_NA24385_son/latest/GRCh38/). For SVs, we compared three linked-read sets (R_9_, R_10_, R_11_) from HG002 with the Tier 1 SV benchmark from Genome in a Bottle [36] and used VaPoR [37] to validate our SV calls based on PacBio CCS reads from NA24385 (Highly-accurate long-read sequencing improves variant detection and assembly of a human genome). We compared SNV and SV calls among the different approaches using vcfeval [38] and truvari [36], respectively.

#### Performance of diploid assembly: influence of total coverage

Diploid assembly by Linked-Reads requires sufficient total read coverage (*C*=*C_R_*×*C_F_*) to generate long contigs and scaffolds. In this experiment, to explore the roles of both physical coverage (*C_F_*) and per-fragment read coverage (*C_R_*), we first generated eight simulated libraries whose total coverage *C* ranged from 16x to 78x: four with *C_R_* fixed and increasing *C_F_* and four with fixed *C_F_*, and increasing *C_R_* (**Table S2**). Contig and scaffold N50s increased along with increasing either *C_F_* or *C_R_* (**Figure 2A** and **2B**). To investigate whether the trend was also present in the real datasets, we analyzed six real libraries (three by varying *C_F_*, and the other three by varying *C_R_*; **Figure 1**): as *C* increased, we varied *C_F_* and *C_R_* independently by fixing the other parameter. Contig and scaffold N50s also increased in these simulation (**Figure 2C and 2D**) and real linked-read sets (**Figure 2E and 2F**) as a function of total coverage C. Contig lengths did increase a little (621.4kb to 758.1kb for simulation; 110.7kb to 119.6kb for real data) when *C* was increased beyond 56X. Accuracy, which we define as the ratio between NA50 (N50 after breaking contigs or scaffolds at assembly errors) and N50 (**Figure 2C** and **2E**), changed 18% for simulation and 7% for real data (587.5kb to 713.3kb for simulation; 97.1kb to 104.5kb for real data). For scaffolds in the real data sets, when *C* increased from 48X (*R*_3_) to 67X (*R*_4_), both scaffold N50 and NA50 were significantly improved (N50: 13.4Mb to 30.6Mb; NA50: 6.3Mb to 12.0Mb), but the accuracy dropped slightly from 46.6% to 39.1%, which indicated that scaffold accuracy may be refractory to extremely high *C* (**Figure 2F**). These results indicated that assembly length and accuracy were comparable over a broad range of *C_F_* and *C_R_* at constant *C*, which implied that assembly quality was mainly determined by *C*.

**Figure 2.**
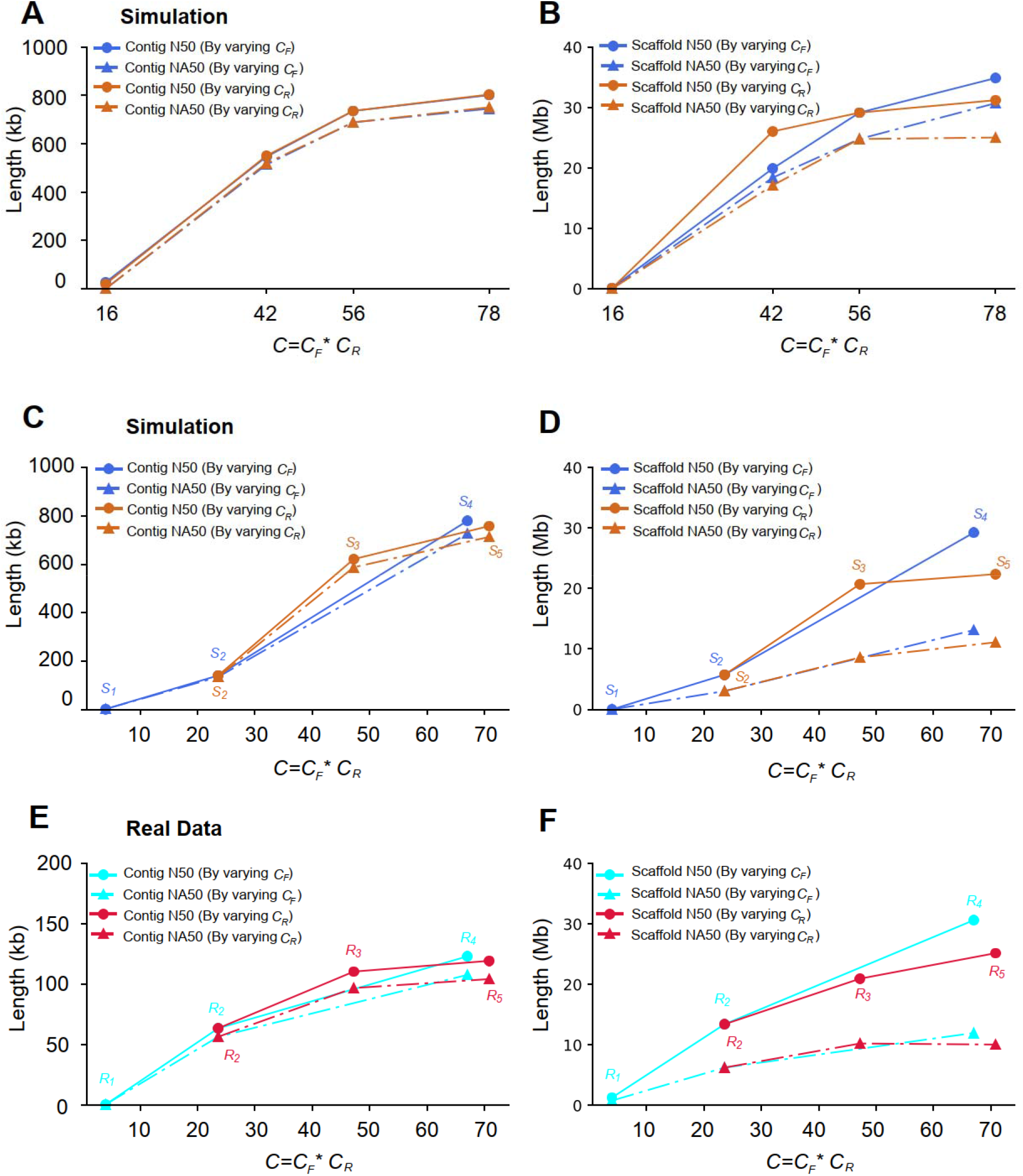
Contig and scaffold lengths (N50 and NA50) as a function of *C_F_* or *C_R_*. **A** and **B**: Simulated Linked-Reads with predefined parameters (**Table S2**); **C** and **D**: Simulated Linked-reads with matched parameters of real Linked-Read data sets (**Figure 1**); **E** and **F**: Real linked-read sets (**Figure 1**).

#### Performance of diploid assembly: influence of fragment length and physical coverage

To investigate if input weighted fragment length (as measured by W*μ_FL_*) influenced assembly quality, we generated four simulated libraries (**Table S3**) with fixed *C_F_* and *C_R_* and a range of fragment lengths (**Figure 3A**). Contig length decreased with increasing fragment length, a trend that was also seen in six real libraries (**Figure 3B**; *C*=56X; *R*_6_ to *R*_11_ in **Figure 1**). We then simulated another six libraries with the same parameters as the real ones to explore the effects of physical coverage at constant *C*=56x (**Figure 3C**). Contig lengths decreased as a function of increasing physical coverage, a trend that is somewhat less clear in real data possibly due to confounding other parameters such as fragment length (**Figure 3D**). The two linked-read sets with the worst contig qualities in NA12878 (*R*_7_) and NA24385 (*R*_10_) also showed a significant increase of the number of breakpoints (**Table S4**)

**Figure 3.**
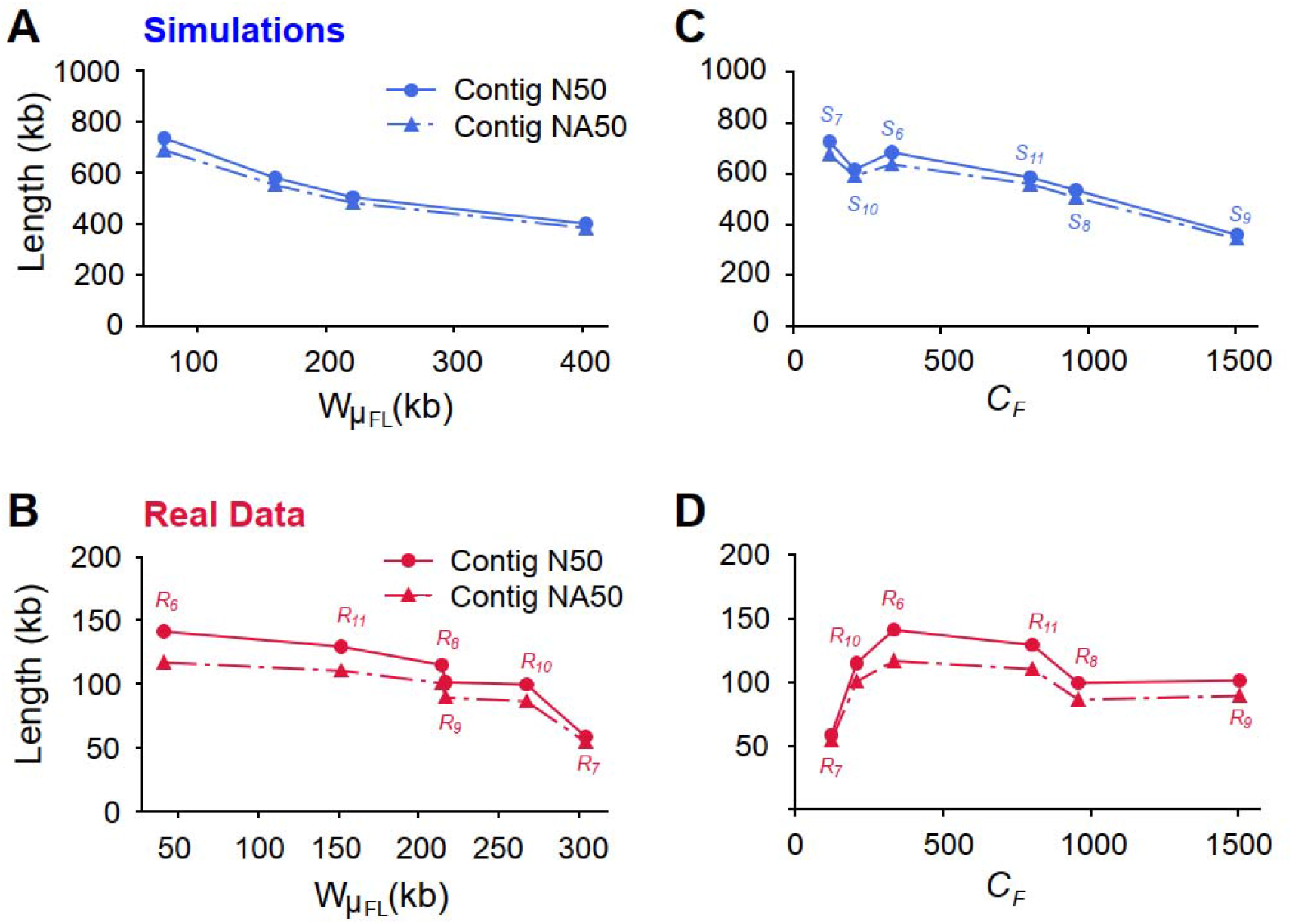
Contig qualities (N50 and NA50) as a function of fragment length *Wμ_FL_* or physical coverage *C_F_*, at *C*=56X. **A** and **C**, results from simulations; **B** and **D**, results from real data.

#### Performance of diploid assembly: nature of the source genome

Assembly errors may occur because of heterozygosity, repetitive sequences, or sequencing error. To illuminate possible sources of assembly error, we performed simulations by generating 10x-like Linked-Reads as above from human chromosome 19, and then quantified assembly error against these synthetic gold standards. Removal of interspersed repeat sequences from the source genome resulted in better contigs with no loss of accuracy in experiments by varying *C_F_, C_R_* and *μ_FL_*(**Figure 4A, 4C** and **4E**) and better scaffolds only if *C_R_* was above 1X (**Figure 4D**). Removal of variation had little effect on contigs and only gave rise to longer scaffolds if *C_R_* was above 0.8X (**Figure S11**), which is difficult to achieve with real libraries. Finally, a 1% uniform sequencing error had no discernible effect (**Figure S12**).

**Figure 4.**
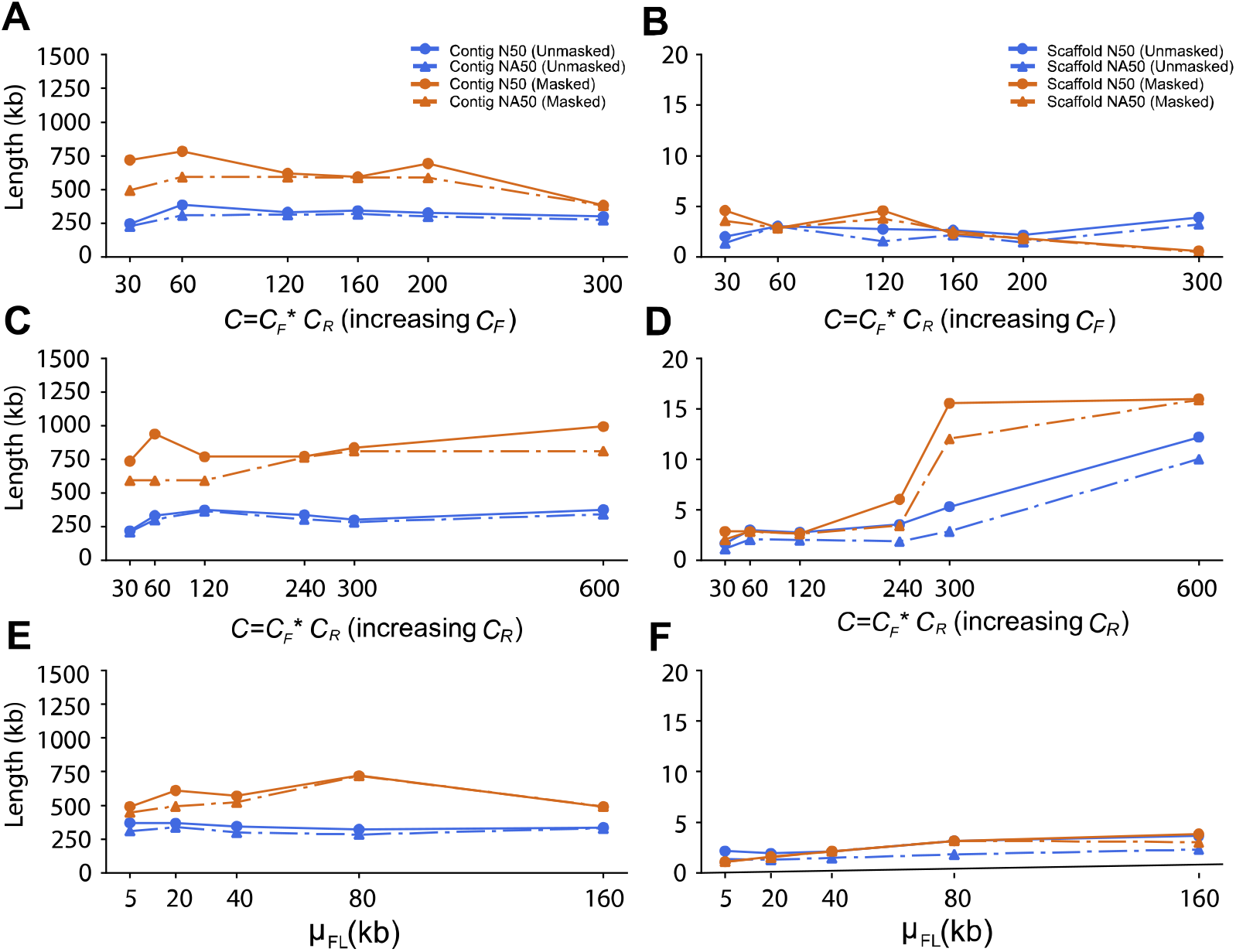
Comparison of contig and scaffold lengths from 10x data with masked and unmasked repetitive sequences by changing *C_F_, C_R_* and *μ_FL_. C_R_* was fixed to 0.2X in **A** and **B**; *C_F_* was fixed to 300X in **C** and **D**; *C_R_* was fixed to 0.2X and *C_F_* was fixed to 300X in **E** and **F**.

#### Performance of diploid assembly: fraction of genome in diploid state

While contiguity is an important parameter for any whole genome assembly, evaluation of diploid assemblies necessitates estimating the fraction of the genome in which the assembly recovered the diploid state. To this end, we divided the contigs generated by Supernova2 into “diploid contigs”, which were extracted from its megabubble structures, and “haploid contigs” from non-megabubble structures. Pairs of scaffolds were extracted as the two haplotypes from megabubble structures if they shared the same start and end nodes in the assembly graph. Diploid contigs were generated by breaking the candidate scaffolds at the sequences with least 10 consecutive ‘N’s and were aligned to human reference genome (hg38) by Minimap2. The genome was split into 500bp windows and diploid regions were defined as the maximum extent of successive windows covered by two contigs, each from one haplotype. Alignment against the human reference genome revealed the overall genome coverages of the six assemblies to be around 91%. For most assemblies, 70%-80% of the genome was covered by two homologous contigs (**Table 1**), with *R*_6_ only reaching 58.9%, probably due to the short fragments of the DNA preparation (*μ_FL_* =24kb). We also analyzed another seven assemblies produced by 10x Genomics, all of which had diploid fractions of about 80% as well (**Table S5**). In the male NA24385, non-pseudoautosomal regions of the X chromosome are hemizygous and should therefore be recovered as haploid regions. Between 79.9% and 87.6% of these regions were covered by one contig exactly depending on the assembled library. Library construction parameters other than fragment length appeared to have had little impact on the proportion of diploid regions (**Tables 1** and **Table S5**).

**Table 1.** Genomic coverage of contigs generated by Supernova2. Non-PAR: non-pseudoautosomal regions of X chromosome. *R*_6_, *R*_7_ and *R*_8_ are female; *R*_9_, *R*_10_ and *R*_11_ are male.

| Linked-reads set | Overall<br>(%) | Diploid<br>regions<br>(%) | Haploid<br>regions<br>(%) | Non-PAR<br>(%) | Total contig<br>length<br>(contig>500bp) | Length of contigs<br>from megabubble<br>(contig>500bp) | Percentage<br>(%) |
| --- | --- | --- | --- | --- | --- | --- | --- |
| $R_6$ | 91.9 | 58.9 | 27.7 | - | 5,632,483,053 | 3,758,345,846 | 66.73 |
| $R_7$ | 91.1 | 73.3 | 11.3 | - | 5,613,140,437 | 4,668,186,478 | 83.17 |
| $R_8$ | 91.7 | 77.2 | 9.2 | - | 5,635,127,471 | 4,896,821,850 | 86.90 |
| $R_9$ | 91.3 | 73.4 | 12.2 | 85.9 | 5,637,615,919 | 4,438,175,621 | 78.72 |
| $R_{10}$ | 91.7 | 79.2 | 5.8 | 79.9 | 5,749,001,471 | 4,793,226,150 | 83.37 |
| $R_{11}$ | 91.7 | 78.1 | 7.9 | 87.6 | 5,677,566,094 | 4,723,083,367 | 83.19 |

Overlapping the diploid regions from the assemblies of the same individual revealed that 50.24% and 67.27% of the genome for NA12878 and NA24385 (**Figure S13**), respectively, were diploid in all the three assemblies. NA12878 was lower because of the low percentage of diploid regions in assembly *R*_6_ (**Table 1**). The overlaps were significantly greater than expected by chance (NA12878: 33.3%, p-value=0.0049; NA24385: 45.4%, p-value=0.0029. Chi square test). These observations were consistent with heterozygous variants being enriched in certain genomic segments, in which two haplotypes were more easily differentiated by Supernova2. Phase block lengths were mainly determined by total coverage *C* and increased in real data with increasing fragment length (**Figure S14, Table S6**).

#### Performance of diploid assembly: quality of variant calls

The ultimate goal of human genome assembly is to accurately identify genomic variants. We compared the SNVs and SVs from our assemblies with the calls from referenced-based processing of standard Illumina and 10x data, and benchmarked them using gold standard from Genome in a Bottle and PacBio CCS reads. We found the SNVs from referenced-based processing of standard Illumina and 10x data were comparable and both of them were better than assembly-based calls (**Table S7 and S8**) For SVs, our assemblies generated many calls that were missed by the reference-based strategy (**Table S9-S12**) and even by the Tier 1 benchmark of Genome in a Bottle (**Table S13**), and half of the deletions and a majority of insertions could be validated by PacBio CCS reads (**Table S14**).

## Discussion

In this study, we investigated human diploid assembly using 10x Linked-Read sequencing data on both simulated and real libraries. We developed the simulator LRTK-SIM to examine the likely impact of parameters in diploid assembly and compared results from simulated reads to those from real libraries. We thus determined the impact of key parameters (*C_R_, C_F_, N_F/P_* and *μ_FL_*/W*μ_FL_*) with respect to assembly continuity and accuracy. Our study provides a general strategy to evaluate assemblies of 10x data and may have implications for the evaluation of other barcode-based sequencing technologies such as CPTv2-seq [39] or stLRF [40] in the future.

### 10x Practicalities

For standard Illumina sequencing, library complexity is usually sufficient to generate tremendous numbers of reads from unique templates and read coverage can be increased simply by sequencing more. However, the 10x Chromium system performs amplification in each partition, and generally only about 20% to 40% of the original long fragment sequence can be captured as short fragments and eventually as reads, resulting in shallow sequencing coverage per fragment. Sequencing more deeply does not increase the per-fragment coverage much as most of the extra reads are from PCR duplicates. The solution is to sequence multiple 10x libraries constructed from the same DNA preparation and merge them for analysis. This means that *C_R_* remains in the standard range where PCR duplicates are relatively rare, but *C_F_* increases proportionally to the number of libraries used. A practical limitation to this approach is that Supernova2 limits the number of barcodes to 4.8 million.

Our results showed that in practice, *C_F_* should be between 335X and 823X, but no larger than 1000X, given the optimal coverage of *C*=56X recommended by 10x and the requirement for sufficient per-fragment read coverage. Surprisingly, we observed that including more extremely long fragments was detrimental for assembly quality. This is possibly due to the loss of barcode specificity for fragments spanning repetitive sequences. From a computational perspective, too many long fragments are harmful to deconvolving the *de bruijn* graph, as more complex paths need to be picked out. In our experiments, W*μ_FL_* between 50kb and 150kb is the best choice to generate reliable assemblies.

### Parameters driving assembly quality

Our results regarding assembly quality, and the 10x parameters that influence it, may be useful for efforts in which *de novo* assemblies are important for generation of an initial reference sequence. We show that maximization of N50 does not necessarily reflect assembly quality, which we were able to compare to NA50 because there exists a high-quality human reference genome. Contig and scaffold lengths mostly increased with ascending sequencing coverage, and at sufficient overall sequence coverage it did not matter much whether the increasing coverage *C* was accomplished by increasing *C_R_* or *C_F_*. However, both contig and scaffold accuracy decreased with increasing C. We also found, counterintuitively, that contig and scaffold length mostly decreased with increasing fragment length, a phenomenon that may be due to the specific implementation; however, until there is another assembler that can be compared to Supernova2 it will not be possible to reason about this effect. In addition, intrinsic properties of the genome matter greatly, as removal of repeats or lack of variation dramatically improves assembly quality.

Diploid assembly is the appropriate approach for assembly of genomes of diploid organisms that harbor variation. Therefore, an important metric to evaluate diploid assembly is the fraction of the genome that is assembled in a diploid state. The short input fragment length of *R*_6_ resulted in roughly 20% less of the genome in a diploid state (<60% vs <80%) compared to the other libraries of the same individual. This observation suggests that in addition to metrics such as N50, evaluation of assembly quality should also include the fraction of the genome (or the assembly) that is in a diploid state.

### Cost-benefit analysis

Overall, we have attempted to give practical guidelines to assembly of 10x data with Supernova2 and evaluate the performance across a wide range of metrics. Arguably, the metric that matters most in the context of a personal genome is the discovery of variation that lower-cost approaches do not enable. We estimate that the cost increase over standard Illumina sequencing is about 2x, given the 10X preparation cost and the higher level of sequence coverage required. There may be many applications for which this combination of excellent single nucleotide variant detection (via barcode-aware read mapping) and precise structural variant discovery (via assembly), achieved by the same data set, is worth the price.

### Comparison with hybrid assemblies

Hybrid assembly strategies have been applied successfully to produce human genome assembly of long contiguity [13, 14, 41]. In these studies, long contigs are first produced by single-molecule long-reads, such as PacBio (NG50=1.1Mb; [13]) or Nanopore (NG50=3.21Mb; [14]) comparing favorably to our best results for Linked-Reads assemblies (NG50=236kb). Scaffolding is then performed with complementary technologies such as BioNano to capture chromosomal level long-range information. It promoted the scaffold N50 of PacBio to 31.1Mb [13] and Illumina mate-pair sequencing with 10x data to 33.5Mb [25]. Using SuperNova2, the scaffold N50 from our studies reached ~27.86Mb (*R*_6_) on the basis of 10x data alone, suggesting that 10x technology gives broadly comparable results at a fraction of the price of long-read-based hybrid assemblies.

## Availability of supporting data

The raw sequencing data are deposited in the Sequence Read Archive and the corresponding BioProject accession number is PRJNA527321. Diploid assemblies and the codes for comparison are currently available at http://mendel.stanford.edu/supplementarydata/zhang_SN2_2019 and https://github.com/zhanglu295/Evaluate_diploid_assembly. LRTK-SIM is publicly available at https://github.com/zhanglu295/LRTK-SIM.

## Additional files

**Table S1.** Parameters of libraries prepared for NA12878 and NA24385.

**Table S2.** Parameters used to generate linked-read sets for evaluating the impact of *C_F_* and *C_R_* on assemblies.

**Table S3.** Parameters used to generate linked-read sets for evaluating the impact of *μ_FL_* and *N_F/P_* on assemblies.

**Table S4.** Contig misassemblies and recovered transcripts of the six assemblies.

**Table S5.** Genomic coverage and fraction of contigs in diploid state generated by Supernova2 for the seven libraries prepared by 10x Genomics. Non-PAR: non-pseudoautosomal regions of X chromosome. WFU, YOR, YORM, PR are female; HGP, ASH and CHI are male.

**Table S6.** Phase block N50s of the six assemblies.

**Table S7.** Comparison SNV calls from standard Illumina data, 10x reference-based calls, and assembly-based calls for NA12878. All calls were compared to the Genome in a Bottle benchmark.

**Table S8.** Comparison SNV calls from standard Illumina data, 10x reference-based calls, and assembly-based calls for NA24385. All calls were compared to the Genome in a Bottle benchmark.

**Table S9.** Comparison of SV calls from standard Illumina data and 10x assembly-based calls for NA12878.

**Table S10.** Comparison of SV calls from standard Illumina data and 10x assembly-based calls for NA24385.

**Table S11.** Comparison of SV calls from 10x reference-based and assembly-based calls for NA12878.

**Table S12.** Comparison of SV calls from 10x reference-based and assembly-based calls for NA24385.

**Table S13.** Comparison of SV calls from our de novo assemblies with the Tier 1 SV benchmark from Genome in a Bottle.

**Table S14.** Proportion of assembly-based SV calls supported by PacBio CCS reads.

**Figure S1. Basic statistics for *L*_1*L*_.** The distributions of **A**. the number of fragments per partition; **B**. sequencing depth per fragment; **C**. probability density function of unweighted fragment lengths; **D**. cumulative density function of unweighted fragment lengths; **E**. reversed cumulative density function of unweighted fragment lengths; **F**. reversed cumulative density function of weighted fragment lengths.

**Figure S2. Basic statistics for *L*_1*M*_.** The distributions of **A.** number of fragments per partition; **B.** sequencing depth per fragment; **C.** probability density function of unweighted fragment lengths; **D.** cumulative density function of unweighted fragment lengths; **E.** reversed cumulative density function of unweighted fragment lengths; **F.** reversed cumulative density function of weighted fragment lengths.

**Figure S3. Basic statistics for *L*_1*H*_.** The distributions of **A**. number of fragments per partition; **B**. sequencing depth per fragment; **C**. probability density function of unweighted fragment lengths; **D**. cumulative density function of unweighted fragment lengths; **E**. reversed cumulative density function of unweighted fragment lengths; **F**. reversed cumulative density function of weighted fragment lengths.

**Figure S4. Basic statistics for *L*_2_.** The distributions of **A.** number of fragments per partition; **B**. sequencing depth per fragment; **C**. probability density function of unweighted fragment lengths; **D**. cumulative density function of unweighted fragment lengths; **E**. reversed cumulative density function of unweighted fragment lengths; **F**. reversed cumulative density function of weighted fragment lengths.

**Figure S5. Basic statistics for *L*_3_.** The distributions of **A**. number of fragments per partition; **B**. sequencing depth per fragment; **C**. probability density function of unweighted fragment lengths; **D**. cumulative density function of unweighted fragment lengths; **E**. reversed cumulative density function of unweighted fragment lengths; **F**. reversed cumulative density function of weighted fragment lengths.

**Figure S6. Basic statistics for *L*_4_.** The distributions of **A**. number of fragments per partition; **B**. sequencing depth per fragment; **C**. probability density function of unweighted fragment lengths; **D**. cumulative density function of unweighted fragment lengths; **E**. reversed cumulative density function of unweighted fragment lengths; **F**. reversed cumulative density function of weighted fragment lengths.

**Figure S7. Basic statistics for *L*_5_.** The distributions of **A**. number of fragments per partition; **B**. sequencing depth per fragment; **C**. probability density function of unweighted fragment lengths; **D**. cumulative density function of unweighted fragment lengths; **E**. reversed cumulative density function of unweighted fragment lengths; **F**. reversed cumulative density function of weighted fragment lengths.

**Figure S8. Basic statistics for *L*_6_.** The distributions of **A**. number of fragments per partition; **B**. sequencing depth per fragment; **C**. probability density function of unweighted fragment lengths; **D**. cumulative density function of unweighted fragment lengths; **E**. reversed cumulative density function of unweighted fragment lengths; **F**. reversed cumulative density function of weighted fragment lengths.

**Figure S9.** The workflow of LRTK-SIM to simulate linked-reads

**Figure S10.** The effect of *N_F/P_* on human diploid assembly of chromosome 19 by Supernova2, where *C* (*C*=60X; *C_F_*=300X and *C_R_*=0.2X) and *μ_FL_* (*μ_FL_*=37kb) are fixed.

**Figure S11.** Comparison of assembly qualities from 10x data with and without single nucleotide variants by changing *C_F_, C_R_* and *μ_FL_. C_R_* was fixed to 0.2X in **A** and **B**; *C_F_* was fixed to 300X in **C** and **D**; *C_R_* was fixed 0.2X and *C_F_* was fixed 300X in **E** and **F**.

**Figure S12.** Comparison of assembly qualities from 10x data with (1% uniform) and without sequencing error by changing *C_F_, C_R_* and *μ_FL_. C_R_* was fixed to 0.2X in **A** and **B**; *C_F_* was fixed to 300X in **C** and **D**; *C_R_* was fixed 0.2X and *C_F_* was fixed 300X in **E** and **F**.

**Figure S13.** Overlaps of diploid regions for the three libraries from the same sample. Diploid regions for NA12878 (**A**) and NA24385 (**B**). The percentages denote the proportion of genome is diploid.

**Figure S14.** Phase block N50s as a function of different parameter combinations. A. simulated linked-reads with predefined parameters (**Table S5**) by changing *C_F_* and *C_R_*; B. simulated linked-reads with matched parameters of real linked-read sets (**Table S2**) by changing *C_F_* and *C_R_*; **C**. real linked-read sets (**Table S2**) by changing *C_F_* and *C_R_*; **D**. simulated linked-read sets (**Table S3**) with different *Wμ_FL_*; **E.** simulated linked-read sets with matched parameters (**Table S3**) with real linked-read sets as *C*=56X; **F**. real linked-read sets with *C*=56X (**Table S3**).

## Author Contributions

AS conceived the study. LZ and XZ wrote LRTK-SIM and performed the analyses. ZMW prepared the genomic DNA and 10x libraries. LZ, XZ, ZMW and AS analyzed the results and wrote the paper. All authors read and approved the final manuscript.

## Acknowledgements

This research was supported by training and research grants from the National Institute of Standards and Technology. We would like to thank Justin Zook, Marc Salit, Alex Bishara, Noah Spies, Nancy Hansen, David Jaffe, and Deanna Church for informative discussions.

