## Supplementary Material for "Assessment of human diploid genome assembly with 10x Linked-Reads data"

**Supplementary Notes**

**LRTK-SIM**

LRTK-SIM was designed to simulate linked-reads generated by the 10X Chromium instrument and Illumina sequencer, which consists of four key steps:

1. **Generate** **diploid reference genome of NA12878.** (See Methods in main manuscript.)
2. **Simulate long DNA fragments.** Assuming the DNA fragment physical coverage was *C_F_*, the total volume of DNA was represented as *V*=*C_F_***L*, where *L* represented the length of reference genome. Equal physical coverage was assumed for two haplotypes, i.e. $\frac{C_{F}}{2}$. The start point for each fragment was randomly picked across the reference and its unweighted length is generated from an exponential distribution with mean of $\mu_{FL}$.
3. **DNA fragment barcoding.** About 4.79 million 16-mer barcode sequences were available in the Long Ranger whitelist. LRTK-SIM generated ‘pseudo’ partitions with unique barcodes and allocated long DNA fragments to them. The number of fragments per partition was sampled from a Poisson distribution with mean of *N_F/P_*.
4. **Simulate Illumina short reads.** LRTK-SIM assumed short reads cover the long DNA fragments uniformly. The number of reads to be generated were calculated as $\frac{C_{R}\times V}{R_{L}}$, where *R_L_* is the length for short reads (150bp by default). The insert size for paired-end reads follows a normal distribution with mean of 400bp. The empirical base quality was learned from linked-reads that were sequenced by Illumina HiSeq X and the positions of sequencing errors were selected randomly.

LRTK-SIM has two advantages: 1. LRTK-SIM is an all-in-one package and implemented in python; it does not rely on any external packages or programs. 2. It explicitly imitates the generation of linked-reads and includes four key parameters: *C_F_*, *C_R_*, $\mu_{FL}$ and *N_F/P_* to fine-tune modeling of the experimental workflow. LRTK-SIM is speeded up by multithreading and publicly available at <https://github.com/LRTK/LRTK-SIM>.

**Evaluation of diploid assembly**

The performance of assembly was evaluated by QUAST-LG and our in-house programs in github (<https://github.com/zhanglu295/Evaluate_diploid_assembly>). The scaffolds were broken into contigs when encountering at least 10 consecutive ‘N’s. Minimap2 was applied to align contigs against reference genome and QUAST-LG was used to post-process the alignments. Contigs shorter than 500bp and scaffolds shorter than 1kb were removed from evaluation. Contig-aligned blocks were generated by breaking contigs at misassemblies, including relocation, inversion and translocation (according to QUAST-LG).

Similar to contigs, for calculating scaffold NA50 scaffold-aligned blocks were obtained by breaking scaffolds at contig misjoins. For example, if there are three contigs ordered as {*a*, *b*, *c*} in the scaffold, misjoins were classified into four types: 1) Relocation, if the alignment-based order of the contigs was different but on the same chromosome; for example if the contigs were aligned in the order {*a*, *c*, *b*} to the hg38 assembly. 2) Inversion, if two connected contigs were in the same order but one of them had reversed orientation in the alignment. 3) Translocation, if two sequences from different chromosomes were merged into adjacent contigs. 4) Indel, if an internal contig such as *b* was unaligned or removed due to insufficient contig length (<500bp). Indels were not considered misjoins if *b* was shorter than 200bp or the gap between *a* and *c* was within 1000bp of *b*’s length.

**Supplementary Tables**

| **Raw DNA Preparation** | **Sequenced**  **Library** | **Sample id** | **Raw coverage (X)** | $\boldsymbol{\mu}_{\boldsymbol{FL}}$**/W**$\boldsymbol{\mu}_{\boldsymbol{FL}}$  **(kb)** | **PCR duplication (%)** | ***C_F_***  **(X)** | ***C_R_* (X)** |
| --- | --- | --- | --- | --- | --- | --- | --- |
| ${Prep}_{1}$ | $L_{1L}$ | NA12878 | 94 | 21.6/38.7 | 53.08 | 19.3 | 0.83 |
| ${Prep}_{1}$ | $L_{1M}$ | NA12878 | 175 | 22.4/39.7 | 29.24 | 117.6 | 0.54 |
| ${Prep}_{1}$ | $L_{1H}$ | NA12878 | 192 | 24.0/41.1 | 10.92 | 334 | 0.27 |
| ${Prep}_{2}$ | $L_{2}$ | NA12878 | 103 | 79.0/304.3 | 19.97 | 123.2 | 0.41 |
| ${Prep}_{3}$ | $L_{3}$ | NA12878 | 106 | 99.2/214.5 | 11.09 | 958.7 | 0.07 |
| ${Prep}_{4}$ | $L_{4}$ | NA24385 | 117 | 92.1/216.9 | 10.88 | 1504.6 | 0.05 |
| ${Prep}_{5}$ | $L_{5}$ | NA24385 | 100 | 120.8/267.4 | 18.51 | 208.4 | 0.25 |
| ${Prep}_{6}$ | $L_{6}$ | NA24385 | 100 | 64.2/151.7 | 12.39 | 803.3 | 0.08 |

**Table S1.** Parameters of libraries prepared for NA12878 and NA24385.

| **Parameters** | **Liked-Read set** | $\boldsymbol{\mu}_{\boldsymbol{FL}}\boldsymbol{/W}\boldsymbol{\mu}_{\boldsymbol{FL}}$  **(kb)** | ***N_F/P_*** | ***C_F_***  **(X)** | ***C_R_***  **(X)** | ***C***  **(X)** |
| --- | --- | --- | --- | --- | --- | --- |
| *C_F_* | *C_F1_* | 37/75 | 10 | 156 | 0.10 | 16 |
|  | *C_F2_* | 37/75 | 10 | 156 | 0.27 | 42 |
|  | *C_F3_* | 37/75 | 10 | 156 | 0.36 | 56 |
|  | *C_F4_* | 37/75 | 10 | 156 | 0.5 | 78 |
| *C_R_* | *C_R1_* | 37/75 | 10 | 44 | 0.36 | 16 |
|  | *C_R2_* | 37/75 | 10 | 117 | 0.36 | 42 |
|  | *C_R3_* | 37/75 | 10 | 156 | 0.36 | 56 |
|  | *C_R4_* | 37/75 | 10 | 217 | 0.26 | 78 |

**Table S2.** Parameters used to generate linked-read sets for evaluating the impact of *C_F_* and *C_R_* on assemblies.

| **Parameters** | **Liked-Read set** | $\boldsymbol{\mu}_{\boldsymbol{FL}}\boldsymbol{/W}\boldsymbol{\mu}_{\boldsymbol{FL}}$  **(kb)** | ***N_F/P_*** | ***C_F_***  **(X)** | ***C_R_* (X)** | ***C***  **(X)** |
| --- | --- | --- | --- | --- | --- | --- |
| $\mu_{FL}$ | $\mu_{FL1}/W\mu_{FL1}$ | 37/75 | 10 | 156 | 0.36 | 56 |
|  | $\mu_{FL2}/W\mu_{FL2}$ | 80/161 | 10 | 156 | 0.36 | 56 |
|  | $\mu_{FL3}/W\mu_{FL3}$ | 110/221 | 10 | 156 | 0.36 | 56 |
|  | $\mu_{FL4}/W\mu_{FL4}$ | 200/402 | 10 | 156 | 0.36 | 56 |
| *N_F/P_* | *N_F/P1_* | 37/75 | 1 | 156 | 0.36 | 56 |
|  | *N_F/P2_* | 37/75 | 2 | 156 | 0.36 | 56 |
|  | *N_F/P3_* | 37/75 | 4 | 156 | 0.36 | 56 |
|  | *N_F/P4_* | 37/75 | 8 | 156 | 0.36 | 56 |
|  | *N_F/P5_* | 37/75 | 16 | 156 | 0.36 | 56 |

**Table S3.** Parameters used to generate linked-read sets for evaluating the impact of $\mu_{FL}$ and *N_F/P_* on assemblies.

| Linked-reads set | $R_{6}$ | $R_{7}$ | $R_{8}$ | $R_{9}$ | $R_{10}$ | $R_{11}$ |
| --- | --- | --- | --- | --- | --- | --- |
| # of breakpoints | 5,171 | 11,479 | 5,637 | 5,682 | 7,967 | 5,157 |
| # of translocations | 807  (15.6%) | 1,776  (15.4%) | 817  (14.5%) | 956  (16.8%) | 1,742  (21.9%) | 847  (16.4%) |
| # of inversions | 192  (3.7%) | 260  (2.3%) | 185  (3.3%) | 164  (2.9%) | 200  (2.5%) | 206  (4.0%) |
| # of relocations | 4,172  (80.7%) | 9,443  (82.3%) | 4,635  (82.2%) | 4,562  (80.3%) | 6,025  (75.6%) | 4,104  (79.6%) |
| # of misassembled contigs | 4,155 | 9,942 | 4,773 | 4,922 | 6,864 | 4,253 |
| # of fully recovered transcripts | 2,154,830  (89.4%) | 2,118,513  (87.9%) | 2,148,275  (89.2%) | 2,145,800  (89.1%) | 2,155,552  (89.5%) | 2,143,173  (89.0%) |
| # of partially recovered transcripts | 4,194  (0.17%) | 10,383  (0.43%) | 5,721  (0.24%) | 6,730  (0.28%) | 5,266  (0.22%) | 4,953  (0.21%) |

**Table S4.** Contig misassemblies and recovered transcripts of the six assemblies.

| **Library** | **Overall (%)** | **Diploid regions (%)** | **Haploid regions (%)** | **Non-PAR (%)** |
| --- | --- | --- | --- | --- |
| HGP | 91.9% | 79.7% | 4.59% | 88.8% |
| ASH | 91.8% | 79.5% | 5.26% | 88.0% |
| WFU | 91.9% | 76.5% | 8.59% | - |
| CHI | 91.8% | 78.3% | 7.50% | 87.7% |
| YOR | 91.8% | 80.1% | 2.27% | - |
| YORM | 91.9% | 76.7% | 2.68% | - |
| PR | 91.9% | 77.2% | 7.89% | - |

**Table S5.** Genomic coverage and fraction of contigs in diploid state generated by Supernova2 for the seven libraries prepared by 10x Genomics. Non-PAR: non-pseudoautosomal regions of X chromosome. WFU, YOR, YORM, PR are female; HGP, ASH and CHI are male.

| Linked-reads set | $R_{6}$ | $R_{7}$ | $R_{8}$ | $R_{9}$ | $R_{10}$ | $R_{11}$ |
| --- | --- | --- | --- | --- | --- | --- |
| Phase Block N50 | 835kb | 2.90Mb | 3.98Mb | 2.69Mb | 6.18Mb | 4.13Mb |

**Table S6.** Phase block N50s of the six assemblies.

|  | **Illumina** | **Reference**  **(R_6_)** | **Assembly**  **(R_6_)** | **Reference**  **(R_7_)** | **Assembly**  **(R_7_)** | **Reference**  **(R_8_)** | **Assembly**  **(R_8_)** |
| --- | --- | --- | --- | --- | --- | --- | --- |
| Total variant calls | 4,080,453 | 4,089,855 | 2,827,759 | 4,628,315 | 1,888,717 | 4,382,163 | 2,755,103 |
| True Positive | 3,079,871 | 3,068,248 | 2,275,801 | 3,078,412 | 1,522,097 | 3,081,345 | 2,317,321 |
| False Negatives | 5,722 | 17,330 | 809,803 | 7,168 | 1,563,502 | 4,235 | 768,272 |
| False Positives | 995,329 | 1,017,881 | 551,034 | 1,546,123 | 366,086 | 1,296,857 | 436,822 |
| Precision | 0.756 | 0.751 | 0.805 | 0.666 | 0.806 | 0.704 | 0.841 |
| Recall | 0.998 | 0.994 | 0.738 | 0.998 | 0.493 | 0.999 | 0.751 |
| F1 | 0.860 | 0.856 | 0.770 | 0.799 | 0.612 | 0.826 | 0.794 |

**Table S7**. Comparison SNV calls from standard Illumina data, 10x reference-based calls, and assembly-based calls for NA12878. All calls were compared to the Genome in a Bottle benchmark.

|  | **Illumina** | **Reference**  **(R_9_)** | **Assembly**  **(R_9_)** | **Reference**  **(R_10_)** | **Assembly**  **(R_10_)** | **Reference**  **(R_11_)** | **Assembly**  **(R_11_)** |
| --- | --- | --- | --- | --- | --- | --- | --- |
| Total variant calls | 3,992,991 | - | 2,586,214 | 4,341,986 | 2,510,950 | 4,294,246 | 2,842,246 |
| True Positive | 3,067,419 | - | 2,218,467 | 3,070,102 | 2,058,441 | 3,072,019 | 2,422,596 |
| False Negatives | 10,086 | - | 859,043 | 7,403 | 1,019,069 | 5,480 | 654,914 |
| False Positives | 920,114 | - | 366,815 | 1,268,249 | 451,766 | 1,218,555 | 418,603 |
| Precision | 0.770 | - | 0.858 | 0.708 | 0.820 | 0.716 | 0.853 |
| Recall | 0.997 | - | 0.721 | 0.998 | 0.669 | 0.998 | 0.787 |
| F1 | 0.869 | - | 0.784 | 0.828 | 0.737 | 0.834 | 0.819 |

**Table S8**. Comparison SNV calls from standard Illumina data, 10x reference-based calls, and assembly-based calls for NA24385. All calls were compared to the Genome in a Bottle benchmark.

|  | **R_6_** | **R_7_** | **R_8_** |
| --- | --- | --- | --- |
| Total SV calls | 9,575 | 9,255 | 10,028 |
| Overlap | 2,059 | 2,186 | 7,472 |
| Only in Illumina-based | 3,291 | 3,168 | 2,886 |
| Only in Assembly-based | 7,453 | 6,989 | 7,472 |
| Overlap/Assembly-based | 0.216 | 0.238 | 0.248 |
| Overlap/Illumina-based | 0.382 | 0.406 | 0.458 |

**Table S9.** Comparison of SV calls from standard Illumina data and 10x assembly-based calls for NA12878.

|  | **R_9_** | **R_10_** | **R_11_** |
| --- | --- | --- | --- |
| Total SV calls | 9,698 | 10,866 | 10,477 |
| Overlap | 1,568 | 1,568 | 1,705 |
| Only in Illumina-based | 3,779 | 3,778 | 3,646 |
| Only in Assembly-based | 8,002 | 9143 | 8,634 |
| Overlap/Assembly-based | 0.164 | 0.146 | 0.165 |
| Overlap/Illumina-based | 0.291 | 0.291 | 0.316 |

**Table S10.** Comparison of SV calls from standard Illumina data and 10x assembly-based calls for NA24385.

|  | **Assembly**  **(R_6_)** | **Assembly**  **(R_7_)** | **Assembly**  **(R_8_)** |
| --- | --- | --- | --- |
| Total SV calls | 9,575 | 9,255 | 10,028 |
| Overlap | 2,060 | 1,537 | 1,432 |
| Only in Reference-based | 1,914 | 986 | 709 |
| Only in Assembly-based | 7,366 | 7,553 | 8,405 |
| Overlap/Assembly-based | 0.219 | 0.169 | 0.146 |
| Overlap/Reference-based | 0.518 | 0.609 | 0.669 |

**Table S11.** Comparison of SV calls from 10x reference-based and assembly-based calls for NA12878.

|  | **Assembly**  **(R_9_)** | **Assembly**  **(R_10_)** | **Assembly**  **(R_11_)** |
| --- | --- | --- | --- |
| Total SV calls | 9,698 | 10,866 | 10,477 |
| Overlap | - | 2056 | 1792 |
| Only in Reference-based | - | 1111 | 709 |
| Only in Assembly-based | - | 8598 | 8475 |
| Overlap/Assembly-based | - | 0.193 | 0.175 |
| Overlap/Reference-based | - | 0.649 | 0.716 |

**Table S12.** Comparison of SV calls from 10x reference-based and assembly-based calls for NA24385. R_9_ failed to be analyzed by Long Ranger due to its extremely large C_F_.

|  | **Assembly**  **(R_9_)** | **Assembly**  **(R_10_)** | **Assembly**  **(R_11_)** |
| --- | --- | --- | --- |
| Total SV calls (Filtered) | 4,535 | 4,965 | 3,743 |
| True Positive | 2,318 | 2,311 | 2,068 |
| False Negatives | 7,381 | 7,388 | 7,631 |
| False Positives | 2,217 | 2,654 | 1,675 |
| Precision | 0.511 | 0.466 | 0.553 |
| Recall | 0.239 | 0.238 | 0.213 |
| F1 | 0.326 | 0.315 | 0.308 |

**Table S13.** Comparison of SV calls from our de novo assemblies with the Tier 1 SV benchmark from Genome in a Bottle.

|  | $R_{9}$ | $R_{10}$ | $R_{11}$ |
| --- | --- | --- | --- |
| Deletion | 0.48 | 0.43 | 0.50 |
| Insertion | 0.77 | 0.74 | 0.76 |

**Table S14.** Proportion of assembly-based SV calls supported by PacBio CCS reads.

**Supplementary Figures**

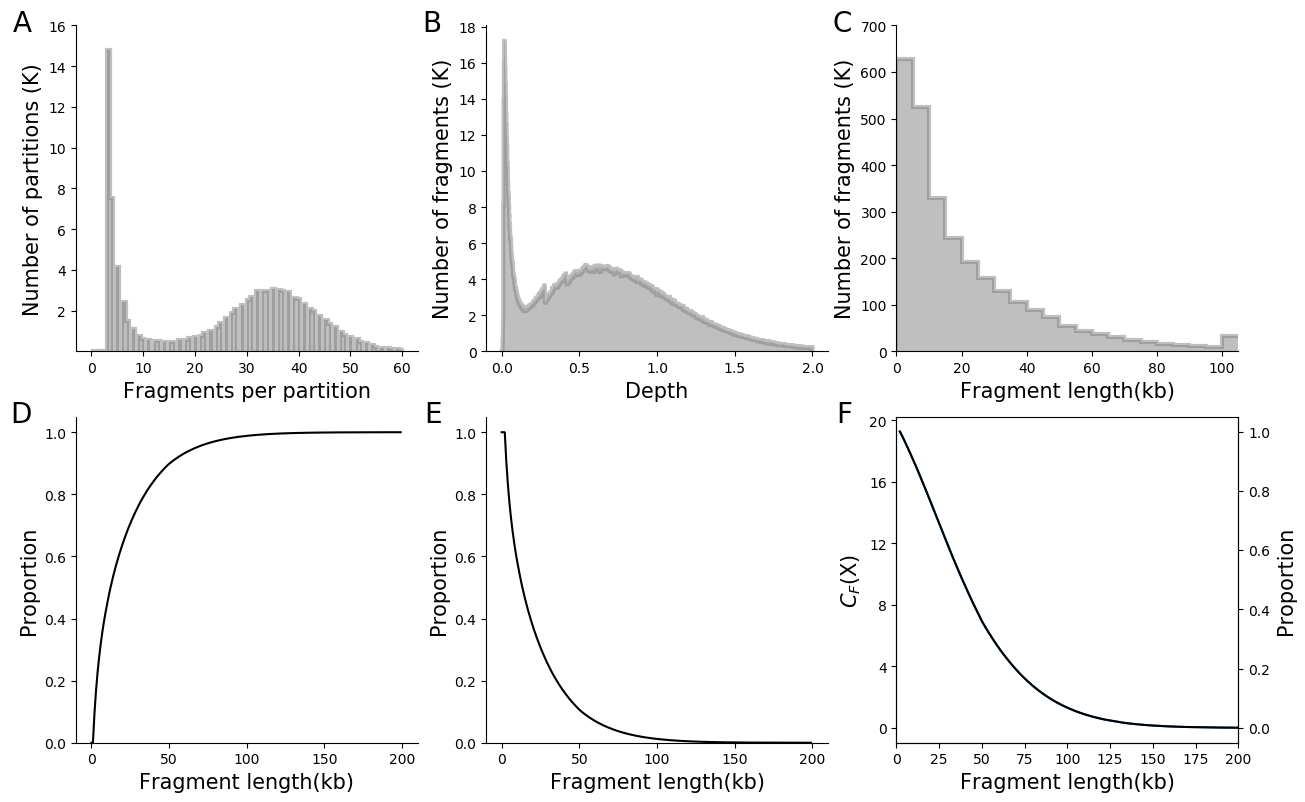

**Figure S1. Basic statistics for** $\boldsymbol{L}_{\mathbf{1}\boldsymbol{L}}$**.** The distributions of **A**. the number of fragments per partition; **B**. sequencing depth per fragment; **C**. probability density function of unweighted fragment lengths; **D**. cumulative density function of unweighted fragment lengths; **E**. reversed cumulative density function of unweighted fragment lengths; **F**. reversed cumulative density function of weighted fragment lengths.

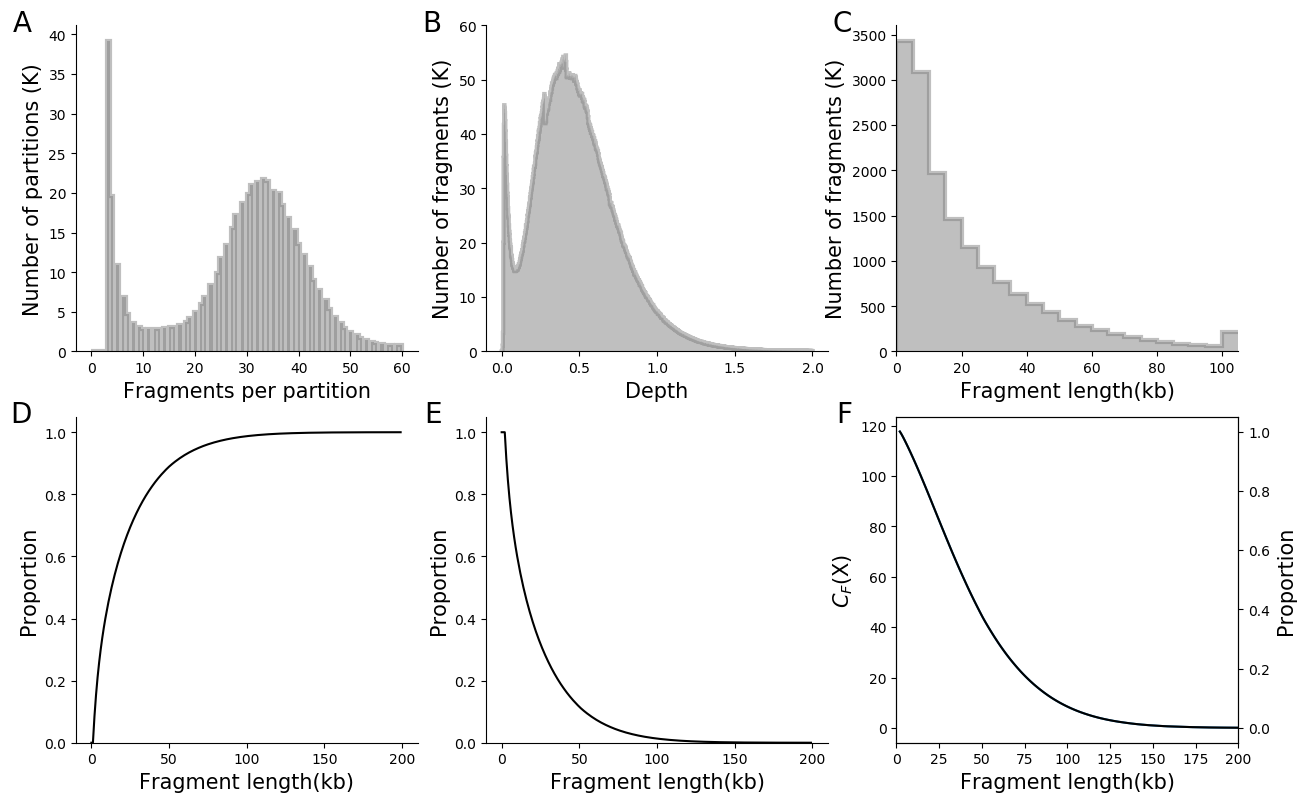

**Figure S2. Basic statistics for** $\boldsymbol{L}_{\mathbf{1}\boldsymbol{M}}$**.** The distributions of **A**. number of fragments per partition; **B**. sequencing depth per fragment; **C**. probability density function of unweighted fragment lengths; **D**. cumulative density function of unweighted fragment lengths; **E**. reversed cumulative density function of unweighted fragment lengths; **F**. reversed cumulative density function of weighted fragment lengths.

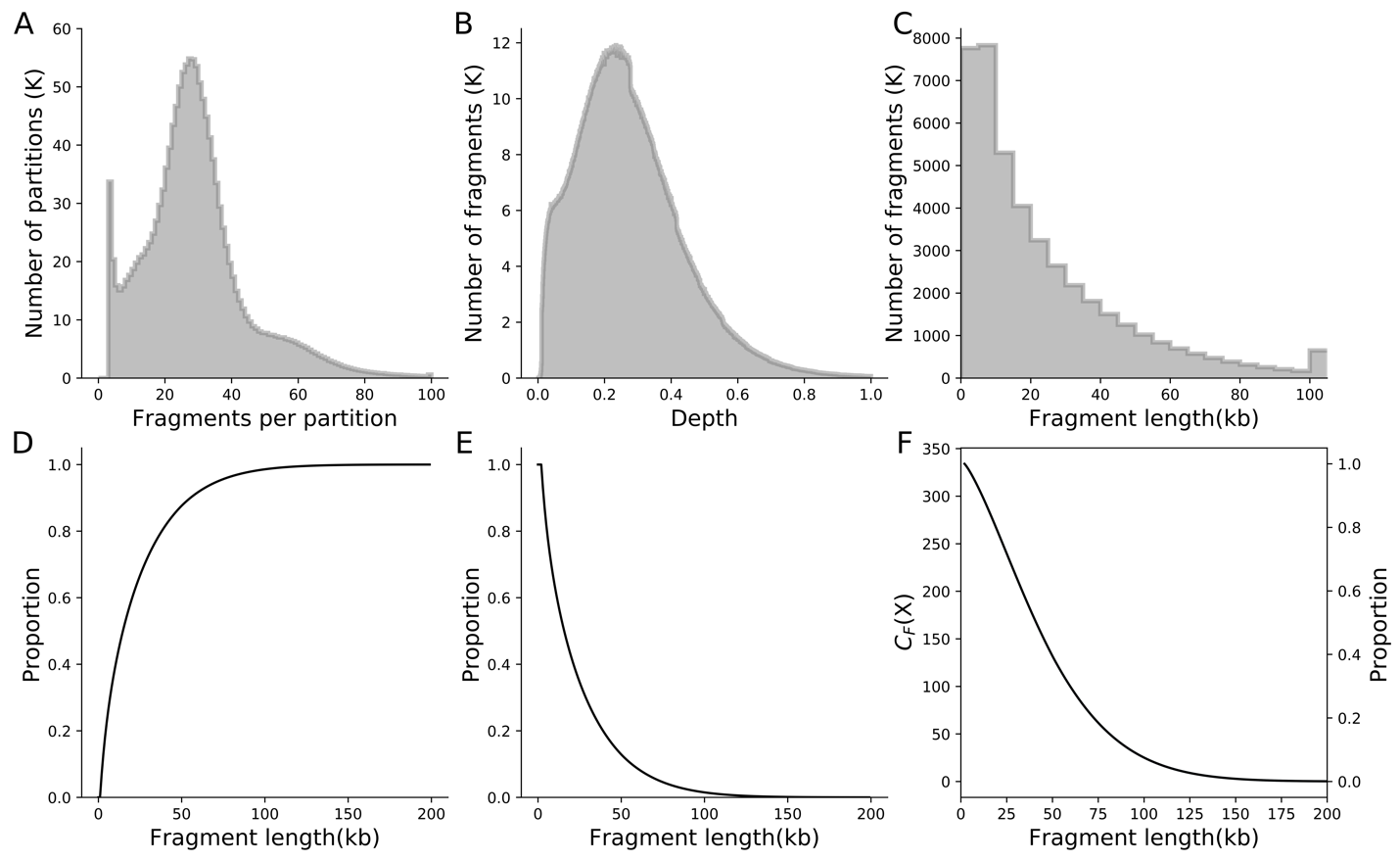

**Figure S3. Basic statistics for** $\boldsymbol{L}_{\mathbf{1}\boldsymbol{H}}$**.** The distributions of **A**. number of fragments per partition; **B**. sequencing depth per fragment; **C**. probability density function of unweighted fragment lengths; **D**. cumulative density function of unweighted fragment lengths; **E**. reversed cumulative density function of unweighted fragment lengths; **F**. reversed cumulative density function of weighted fragment lengths.

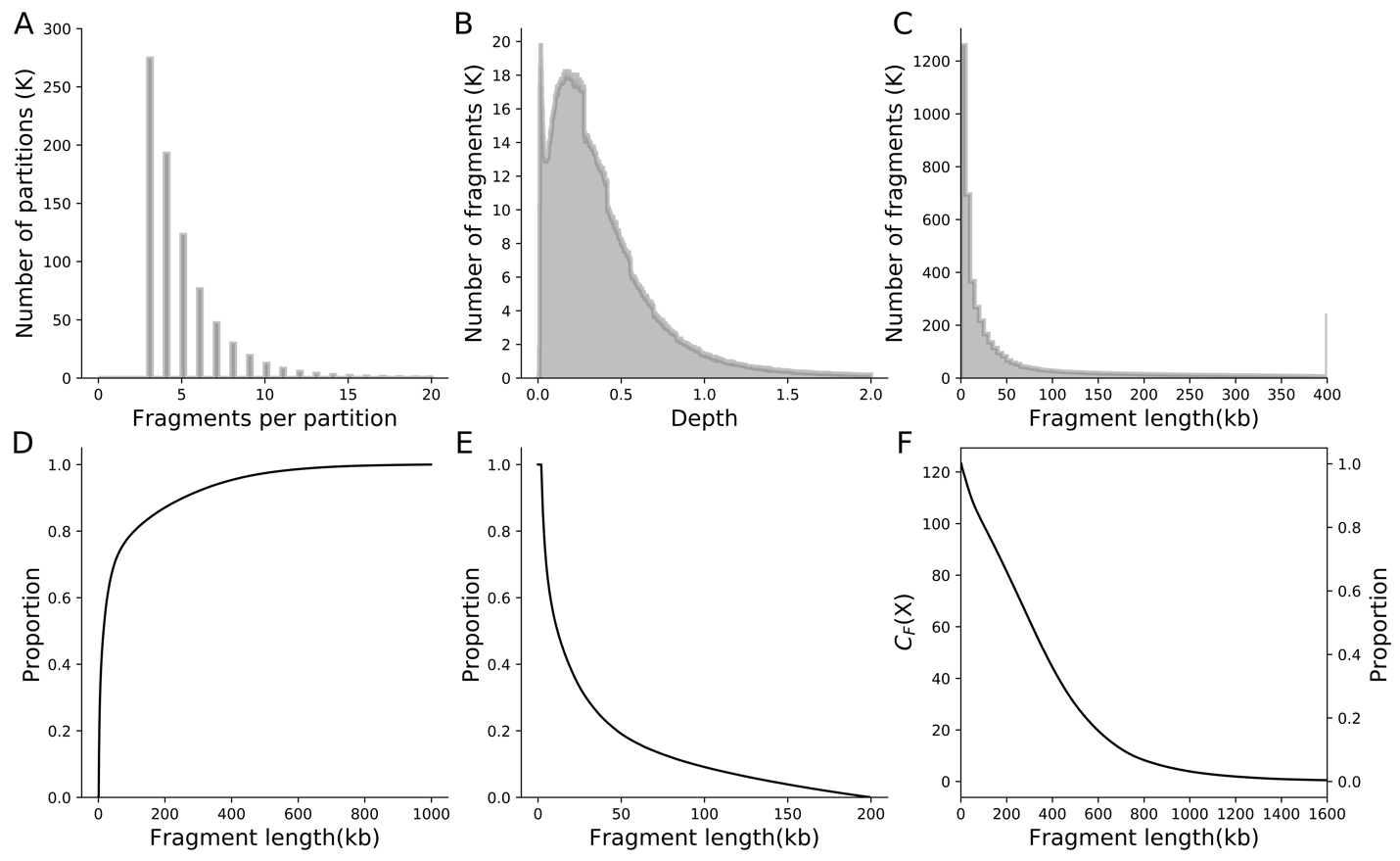

**Figure S4. Basic statistics for** $\boldsymbol{L}_{\mathbf{2}}$**.** The distributions of **A**. number of fragments per partition; **B**. sequencing depth per fragment; **C**. probability density function of unweighted fragment lengths; **D**. cumulative density function of unweighted fragment lengths; **E**. reversed cumulative density function of unweighted fragment lengths; **F**. reversed cumulative density function of weighted fragment lengths.

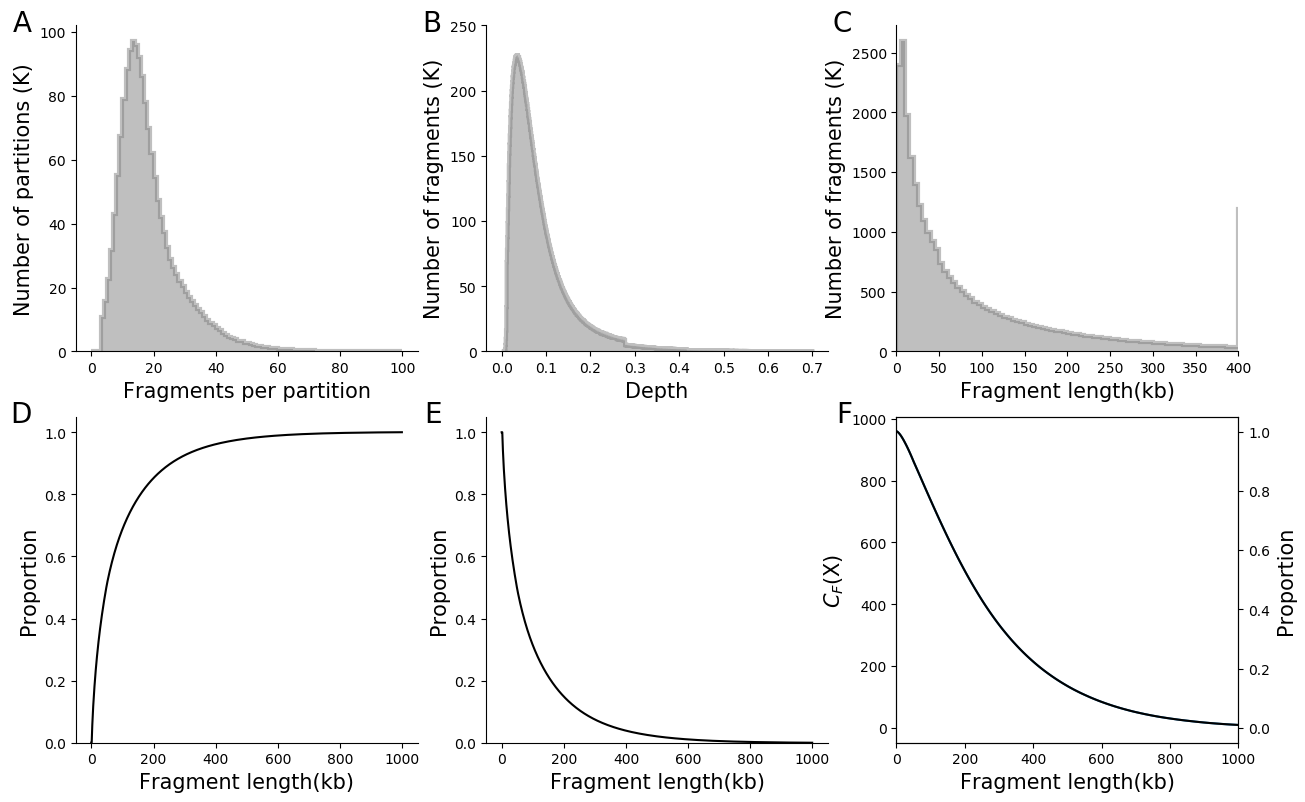

**Figure S5. Basic statistics for** $\boldsymbol{L}_{\mathbf{3}}$**.** The distributions of **A**. number of fragments per partition; **B**. sequencing depth per fragment; **C**. probability density function of unweighted fragment lengths; **D**. cumulative density function of unweighted fragment lengths; **E**. reversed cumulative density function of unweighted fragment lengths; **F**. reversed cumulative density function of weighted fragment lengths.

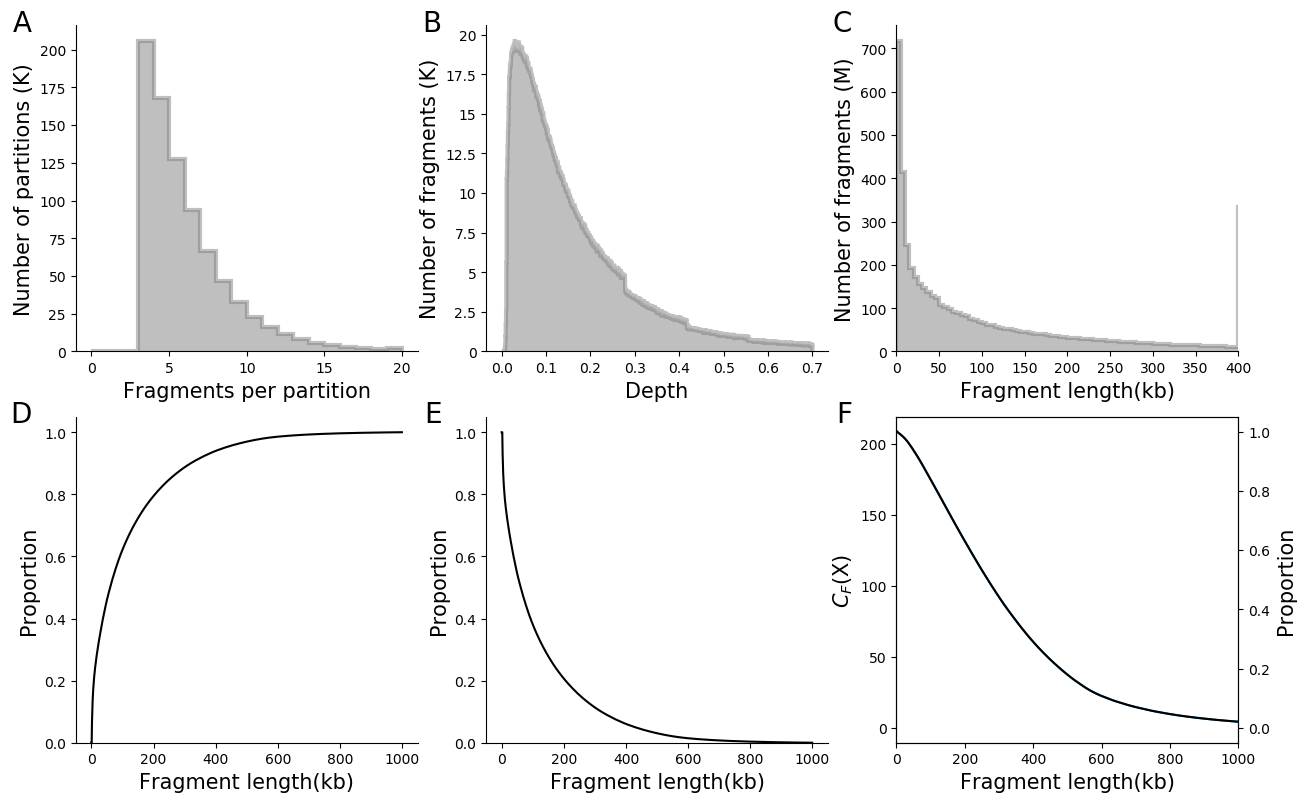

**Figure S6. Basic statistics for** $\boldsymbol{L}_{\mathbf{4}}$**.** The distributions of **A**. number of fragments per partition; **B**. sequencing depth per fragment; **C**. probability density function of unweighted fragment lengths; **D**. cumulative density function of unweighted fragment lengths; **E**. reversed cumulative density function of unweighted fragment lengths; **F**. reversed cumulative density function of weighted fragment lengths.

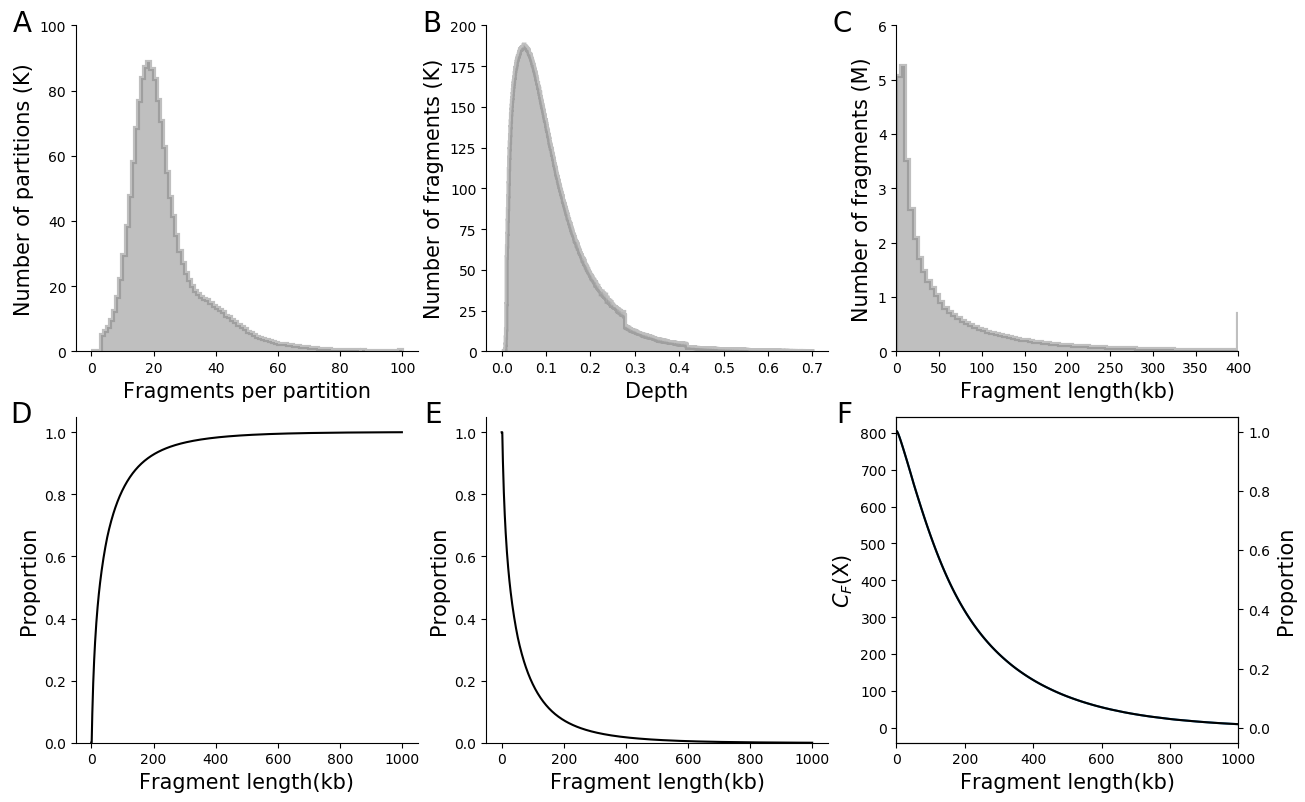

**Figure S7. Basic statistics for** $\boldsymbol{L}_{\mathbf{5}}$**.** The distributions of **A**. number of fragments per partition; **B**. sequencing depth per fragment; **C**. probability density function of unweighted fragment lengths; **D**. cumulative density function of unweighted fragment lengths; **E**. reversed cumulative density function of unweighted fragment lengths; **F**. reversed cumulative density function of weighted fragment lengths.

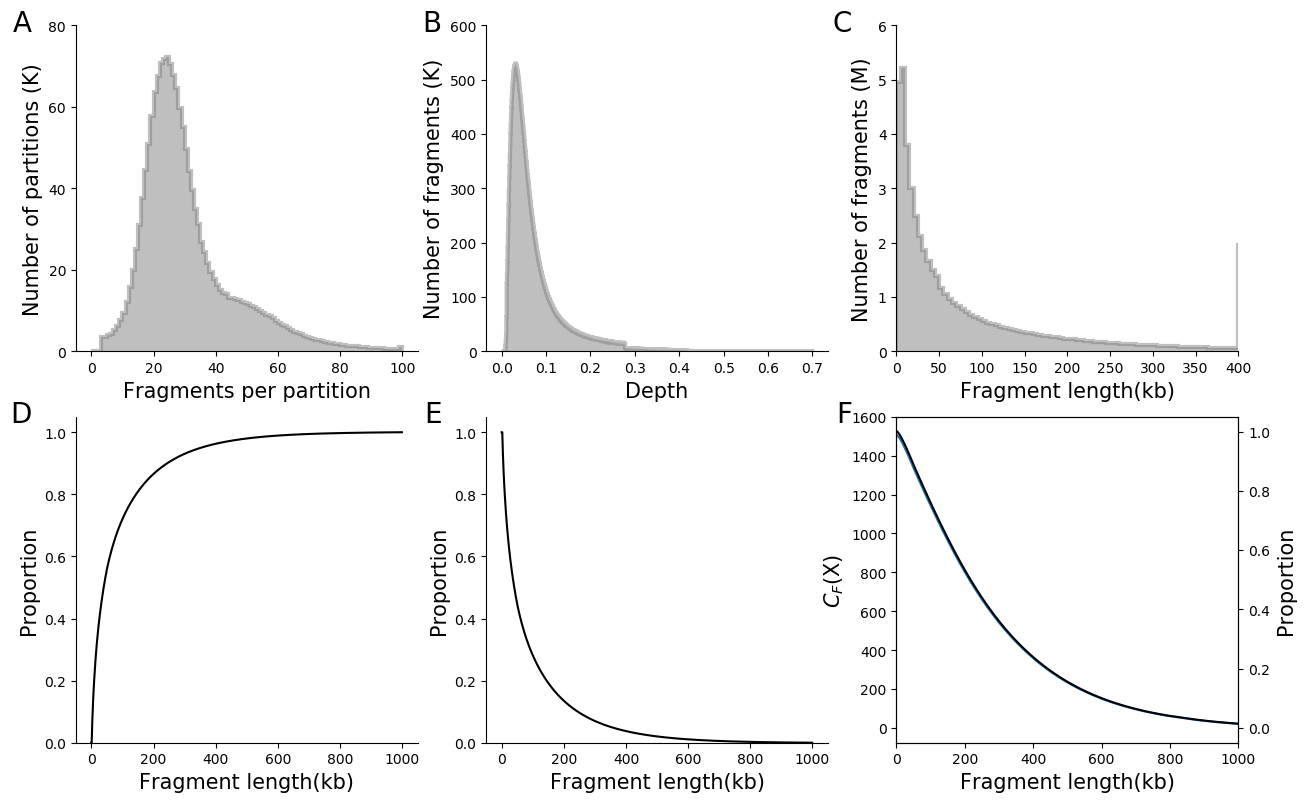

**Figure S8. Basic statistics for** $\boldsymbol{L}_{\mathbf{6}}$**.** The distributions of **A**. number of fragments per partition; **B**. sequencing depth per fragment; **C**. probability density function of unweighted fragment lengths; **D**. cumulative density function of unweighted fragment lengths; **E**. reversed cumulative density function of unweighted fragment lengths; **F**. reversed cumulative density function of weighted fragment lengths.

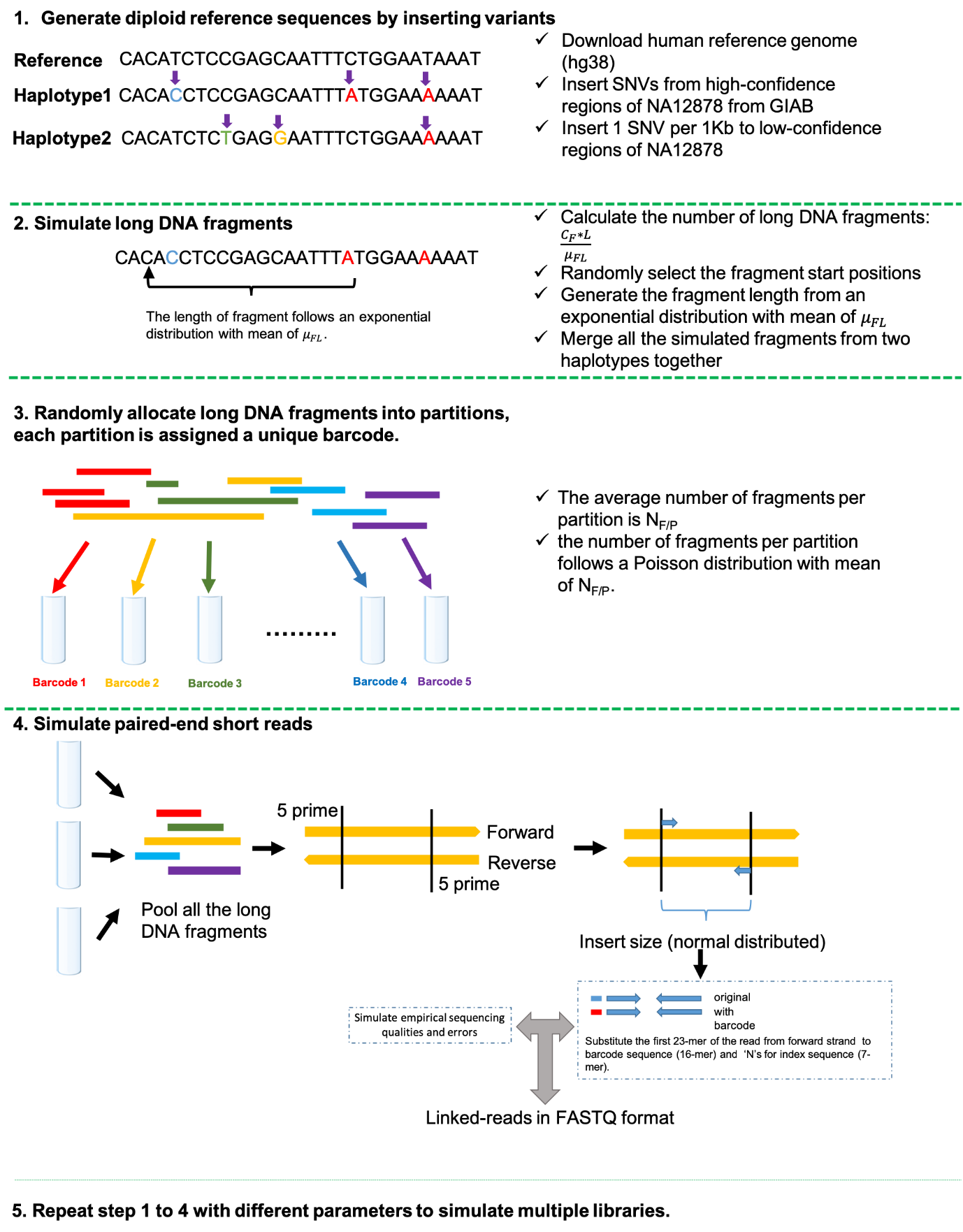

**Figure S9.** The workflow of LRTK-SIM to simulate linked-reads

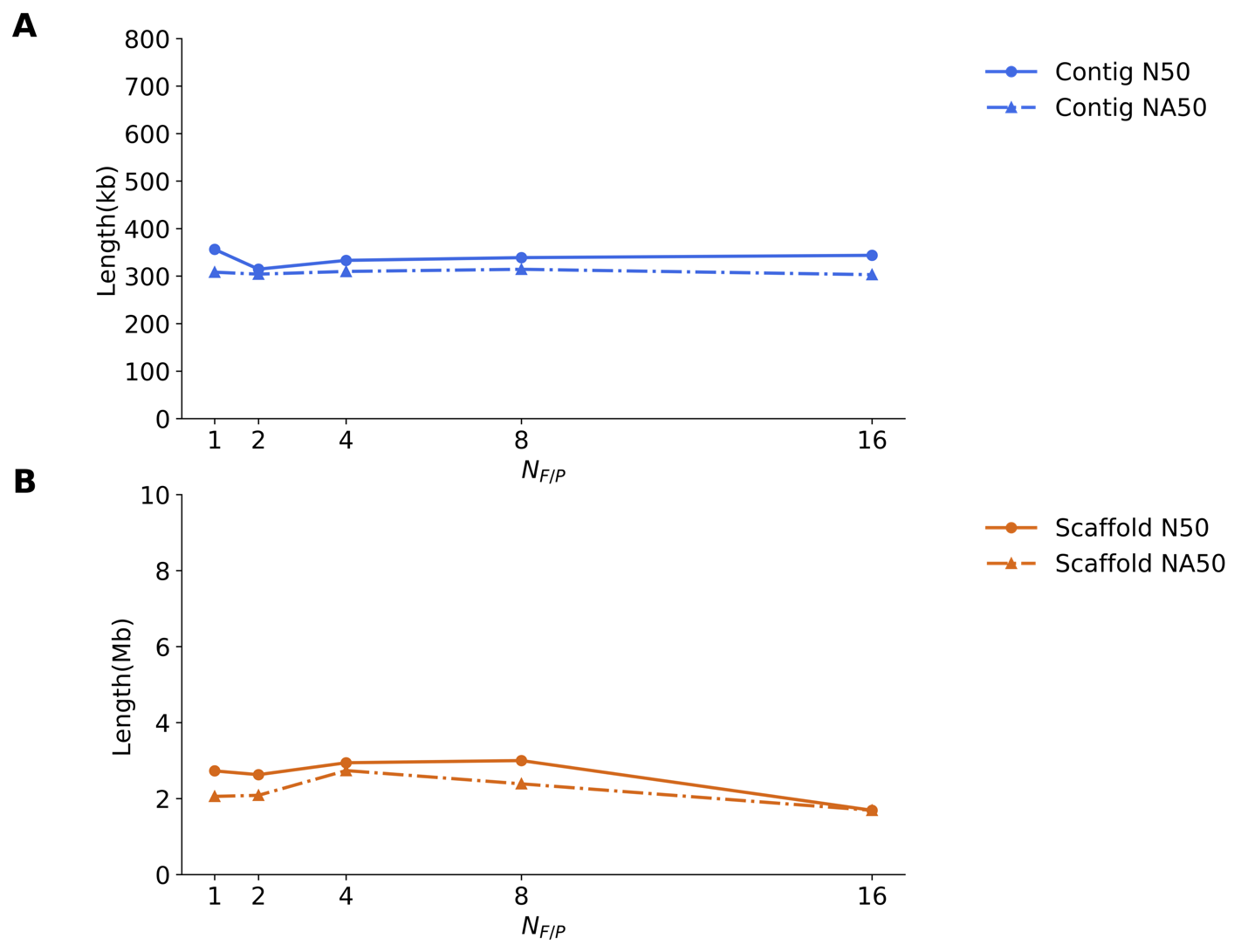

**Figure S10.** The effect of *N_F/P_* on human diploid assembly of chromosome 19 by Supernova2, where *C* (*C*=60X; *C_F_*=300X and *C_R_*=0.2X) and $\mu_{FL}$ ($\mu_{FL}$=37kb) are fixed.

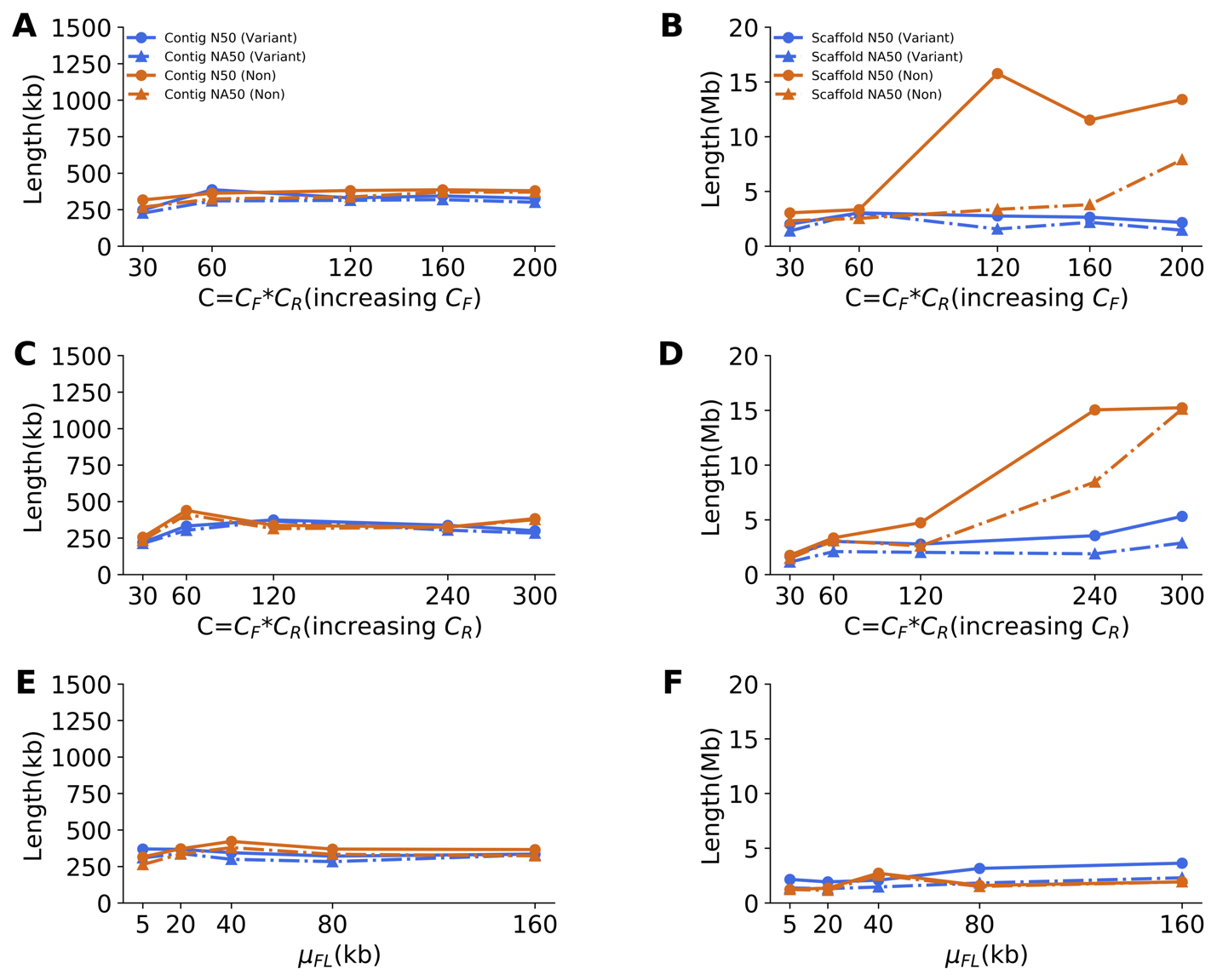

**Figure S11.** Comparison of assembly qualities from 10x data with and without single nucleotide variants by changing *C_F_*, *C_R_* and $\mu_{FL}$. *C_R_* was fixed to 0.2X in **A** and **B**; *C_F_* was fixed to 300X in **C** and **D**; *C_R_* was fixed 0.2X and *C_F_* was fixed 300X in **E** and **F**.

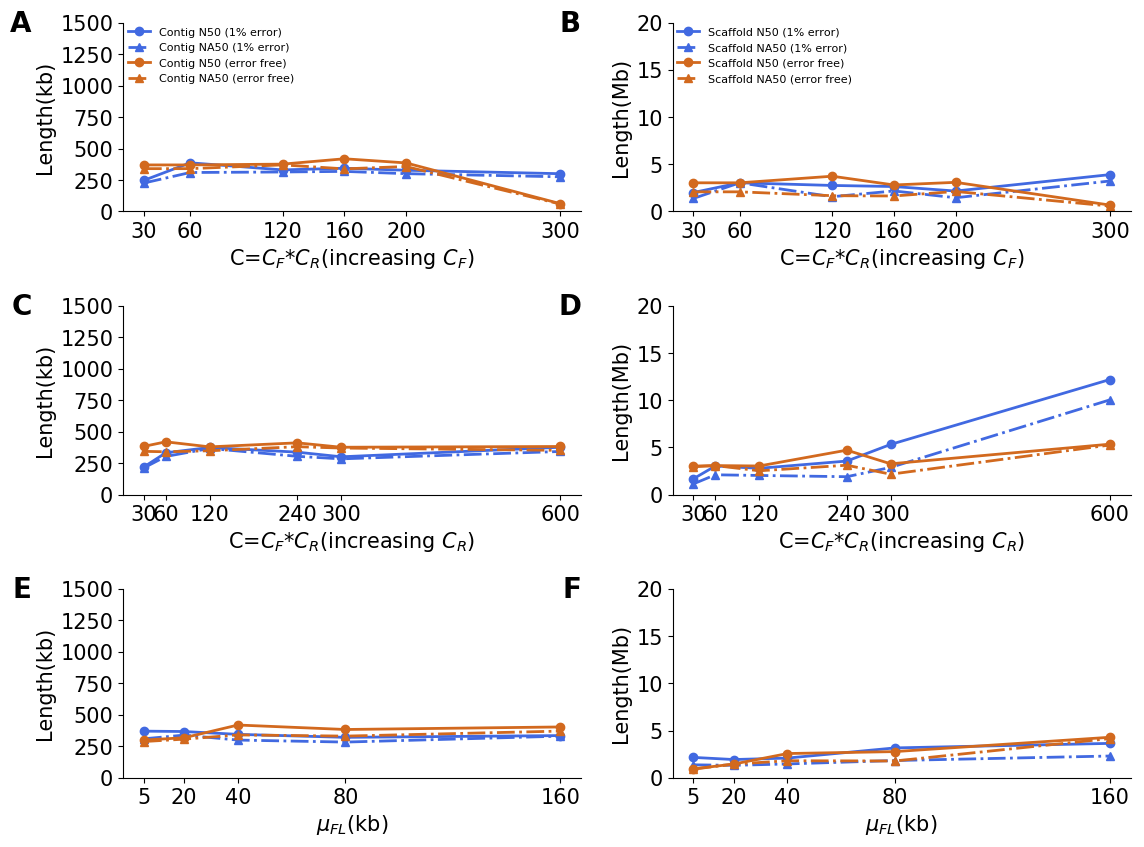

**Figure S12.** Comparison of assembly qualities from 10x data with (1% uniform) and without sequencing error by changing *C_F_*, *C_R_* and $\mu_{FL}$. *C_R_* was fixed to 0.2X in **A** and **B**; *C_F_* was fixed to 300X in **C** and **D**; *C_R_* was fixed 0.2X and *C_F_* was fixed 300X in **E** and **F**.

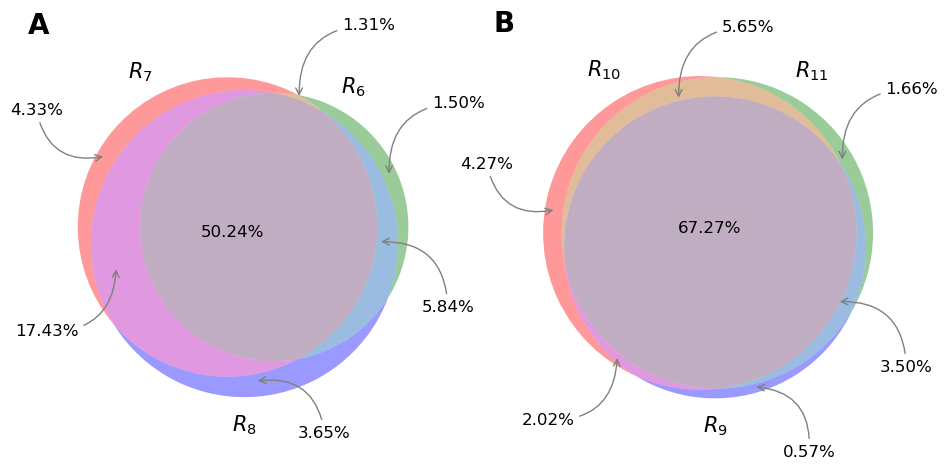

**Figure S13.** Overlaps of diploid regions for the three libraries from the same sample. Diploid regions for NA12878 (**A**) and NA24385 (**B**). The percentages denote the proportion of genome is diploid.

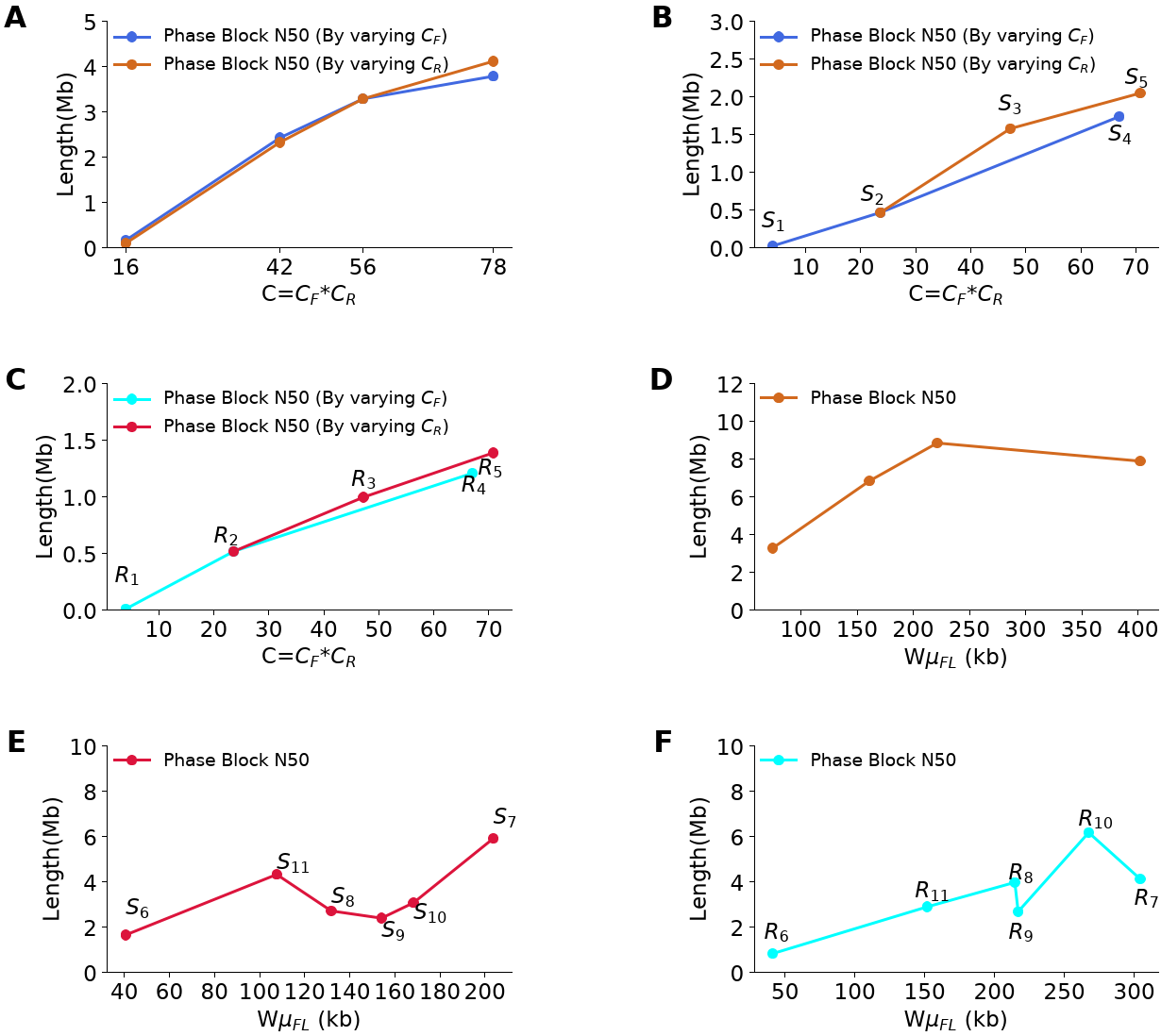

**Figure S14.** Phase block N50s as a function of different parameter combinations. **A**. simulated Linked-Reads with predefined parameters by changing *C_F_* and *C_R_* (**Table S5**); **B**. simulated Linked-Reads with matched parameters of real linked-read sets (**Table S2**) by changing *C_F_* and *C_R_*; **C**. real linked-read sets (**Table S2**) by changing *C_F_* and *C_R_*; **D**. simulated linked-read sets (**Table S3**) with different $W\mu_{FL}$; **E.** simulated linked-read sets with matched parameters (**Table S3**) with real linked-read sets as *C*=56X; **F.** real linked-read sets with *C*=56X (**Table S6**).
